# Leveraging Uncertainty Estimates for Drug Response Prediction in Cancer Cell Lines

**DOI:** 10.64898/2026.04.03.715851

**Authors:** Pascal Iversen, Bernhard Y Renard, Katharina Baum

## Abstract

Machine learning models for drug response prediction in cancer cell lines carry the potential to advance precision oncology by tailoring treatments to the molecular tumor profile. Their application is challenged by variability in prediction quality and distribution shifts between training and application. Uncertainty estimation provides more information on the predictive distribution than point estimates, enabling comprehensive decision support and downstream analysis of predictions. Yet, the most effective uncertainty estimator is domain-specific. In this work, we benchmark uncertainty-aware models for drug response prediction. We focus on epistemic uncertainty via ensemble agreement, and aleatoric uncertainty via distributional modeling, or both. We find that ensemble-based estimates are more sensitive to distribution shift and can flag out-of-distribution examples. In contrast, distributional models yield stronger prediction error reductions among high-confidence subsets. Despite higher computational cost, the combination can provide both advantages: an ensemble of neural networks that estimate a Gaussian predictive distribution can reduce the mean squared error by 64 percent when restricting predictions to the 10 percent most confident drug-cell line pairs, and reliably indicates distribution shifts and platform differences. Beyond benchmarking, probabilistic predictions can identify drugs whose uncertainty bounds overlap with therapeutically relevant ranges. We also show that uncertainty estimates enable a new dimension of model interpretability: by attributing predicted uncertainty to input features, we identify genes that signal unpredictability of drug response rather than sensitivity or resistance. We further demonstrate uncertainty-guided selection of measurements for active learning. In summary, including uncertainty in drug response prediction supports better-informed model application. The code is available at https://github.com/PascalIversen/LUDRP.

## Introduction

Data-driven drug response prediction models aim to identify drug-specific sensitivity or resistance patterns in the omics profile of tumor cell lines ***Barretina et al. (2012***). These models carry the potential to improve therapeutic outcomes by informing preclinical studies or ultimately medical decisions. Large-scale projects such as Genomics of Drug Sensitivity in Cancer (GDSC) screened tumor-derived, cultured cell lines against a wide array of anti-cancer agents ***Yang et al. (2013***), enabling Machine Learning (ML)-based preclinical drug response prediction models. These models can be fine-tuned on data that better represent clinical tumor cases, such as xenografts, and used to identify therapeutic markers using explainable ML, or to recommend drug repurposing candidates ***Manica et al. (2019); Prasse et al. (2022); Lee et al. (2017***). Despite abundant research, challenges persist that hinder their application: Our data-leakage-free, unbiased evaluations show that population-wide model errors are high ***Bernett et al. (2026***). In practice, the prediction error is heteroscedastic, i.e., the prediction quality varies considerably between cell line-drug pairs. Therefore, a naive empirical risk estimate on the test set is a weak estimate for the model’s prediction performance on a specific cell line-drug pair. Further, even if drug response models can be improved to achieve a high average accuracy across the population, they can still show substantial errors for specific cell-line drug pairs ***Chakraborti et al. (2025***).

Additionally, distribution shift (systematic differences between the training and application data) can occur when transferring across screening platforms or to more complex preclinical systems ***Haibe-Kains et al. (2013***). This can lead to a silent failure of applied drug response prediction models. The cell line panels may not adequately represent the diversity of the target application, or measurement conditions might change inadvertently ***Raghavan (2022***). Such shifts have caused failures in applied biomedical ML ***Zech et al. (2018); Wong et al. (2021); Obermeyer et al. (2019***), which motivates us to characterize model robustness early in development. Explicitly modeling uncertainty can ensure that ML-based cancer drug sensitivity models become more useful and reliable. Uncertainty can indicate when to trust a prediction or when additional validation is needed. Further, uncertainty estimates allow a model to restrict its prediction on high-confidence drugs: It may be more important to find one highly effective candidate drug which is predicted with a high confidence, instead of accurately representing the drug response of all available compounds. Conversely, uncertainty enables probabilistic comparisons between predictions. A drug with a moderately unfavorable predicted mean response but high uncertainty may place more probability mass in relevant regions than one with a better mean response but a narrow uncertainty band (see Figure 1 for an example).

**Figure 1.**
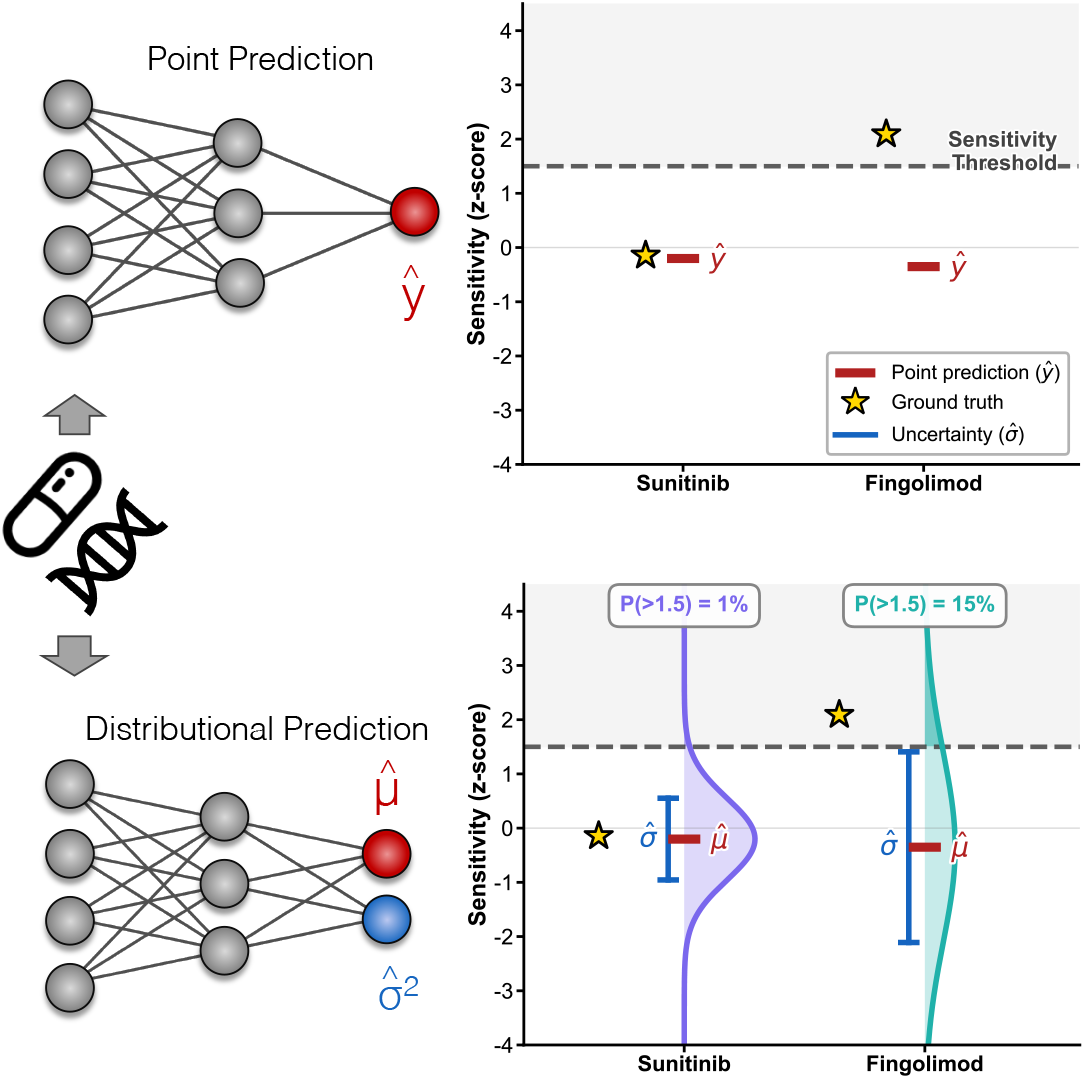
Uncertainty estimates enable more informative drug candidate analyses. Predicted means versus predicted distributions from a Gaussian Neural Network Ensemble for two drugs in the leukemia cell line CESS. Fingolimod has a lower mean (point) prediction but higher uncertainty, placing more probability mass above a drug-wise scaled relative sensitivity threshold at z=1.5, i.e., more sensitive than approximately 93% of cell lines for that drug. Responses (sensitivities, -log IC50) are z-scored per drug. Stars: ground truth relative drug sensitivity. Example selected for illustration. For more predictions for this cell line, see Figure A1.

Uncertainty quantification is an active area of research. Approaches range from Bayesian methods and frequentist inference to evaluating the agreement of ensembles. It is not *a priori* known which method is superior for a given dataset, as they differ in their assumptions, computational cost, and suitability to the model and data at hand ***Abdar et al. (2021); Fakour et al. (2024***). However, comprehensive benchmarks that jointly assess predictive accuracy and the reliability of uncertainty estimates for drug response prediction are lacking. To address this, we here benchmark various uncertainty-aware ML-approaches along these two axes and discuss strategies for their joint evaluation. Uncertainty estimation also opens up downstream applications that we showcase, such as detecting transcriptomic drivers of uncertainty, i.e., genes whose expression is associated with unpredictability of the drug response rather than sensitivity or resistance. These genes may characterize biological states that lead to variable drug responses and would be missed in standard explainability analyses. We further demonstrate that uncertainty estimates can guide active learning to propose informative measurements.

### Related work

Extensive research has been dedicated to identify the best point predictors for drug response ***Adam et al. (2020***), but the community has given little attention to leveraging uncertainty quantification methods in this domain ***Lenhof et al. (2024a). Fang et al. (2018***) and ***Nolte et al. (2024***) propose quantile and deep regression forests, respectively, for constructing prediction intervals in drug response prediction. ***Lenhof et al. (2024b***) focus on applying conformal prediction to achieve reliable drug sensitivity estimation and drug ranking under a novel viability measure. They employ quantile regression forests to estimate the uncertainty in drug-sensitivity regression. ***Rolli et al. (2026***) extend this approach with a cluster analysis that scores how far test samples lie from their nearest training samples to assess the limits of the model applicability. ***Peres da Silva et al. (2021***) weight task- and domain-level aleatoric and epistemic uncertainty to improve transfer from panels to *in vivo* settings. However, these works do not compare the effectiveness of uncertainty quantification methods for drug response prediction. Further, other methods for uncertainty-aware modeling such as distributional neural networks ***Nix and Weigend (1994); Bishop (1994); Koenker and Bassett (1978); Padilla et al. (2022***), Bayesian inference ***MacKay (1992); Gal and Ghahramani (2016***), evidential deep learning ***Amini et al. (2020***), or ensembles of neural networks ***Lakshminarayanan et al. (2017***) have not been considered for drug response prediction. In related molecular property prediction tasks, benchmarks have found optimal uncertainty quantification methods to be highly dataset-, split-, and task-specific, underscoring the need for domain-specific systematic benchmarking ***Greenman et al. (2025); Hirschfeld et al. (2020***).

## Methods

In this work, we compare methods for quantifying uncertainty in drug response prediction and apply these estimates to downstream analytical tasks. First, we evaluate seven uncertainty-aware drug response models on point prediction accuracy and uncertainty quality. Second, we test the ability of these uncertainty estimates to detect out-of-distribution inputs under dataset shifts and generalization settings. Third, we leverage the generated uncertainty estimates for model interpretation and active learning. The code for all experiments, analyses, and figures is available at https://github.com/PascalIversen/LUDRP. The input data to run the code is available at https://zenodo.org/records/19219090.

### Data

We train and evaluate the models on logarithmic IC50 drug response data from the publicly available GDSC dataset. We z-scale the log IC50 values over all drug-cell line pairs for each fold using statistics calculated on the train splits. We use mRNA expression data as cell line features, which are hypothesized to have the highest predictive power ***Manica et al. (2019***). Following Manica et al., we use expression from 2019 genes selected by network diffusion from the STRING protein-protein interaction network around drug targets. We process GDSC’s microarray mRNA expression data using an arcsinh transformation, followed by gene-wise standardization (z-scoring) based on statistics computed from the training set ***Manica et al. (2019***). We use RDKit^1^ to encode the chemical structure of the anti-cancer drugs as 128-bit Morgan fingerprints ***Rogers and Hahn (2010***). After filtering cell lines and drugs with missing gene expression values or drug fingerprints, respectively, we are left with 924 cell lines, 367 drugs, and 292,165 IC50 values. To test whether our main findings are specific to GDSC and for cross-dataset experiments, we additionally reproduce the benchmark findings on the independent Cancer Therapeutics Response Portal (CTRPv2) ***Seashore-Ludlow et al. (2015***) screen with Broad’s CCLE RNA-seq mRNA expression data ***Ghandi et al. (2019***), which after the same filtering comprises 823 cell lines, 466 drugs, and 180,490 logarithmic IC50 values. We further use BeatAML ***Tyner et al. (2018***) for cross-dataset experiments, a dataset of *ex vivo* drug responses in primary samples from acute myeloid leukemia (AML) patients. After restricting to the 80 drugs it shares with CTRPv2 and to samples with available RNA-seq gene expression, it contains 476 samples and 10,017 IC50 values.

### Benchmark

#### Uncertainty in Drug Response Prediction

Commonly, drug response prediction is considered as a regression task ***Adam et al. (2020***): Given a set of cell lines and drugs indexed by *i* = 1, …, *N*_*c*_ and *j* = 1, …, *N*_*d*_, respectively, we observe drug response values *y*_*ij*_ ∈ ℝ (e.g., log IC50) for a subset of samples (*i, j*) ∈ *O*, where *O* ⊆ {1, …, *N*_*c*_} × {1, …, *N*_*d*_ } denotes the index set of observed cell line–drug pairs. We define 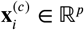 as the feature vector of cell line *i* and 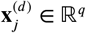 as the feature vector of drug *j*. The training data consists of tuples

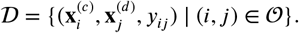

The goal is to learn a predictive function *f* ∶ ℝ^*p*^ × ℝ^*q*^ → ℝ such that for any 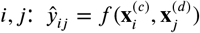 approximates the true drug response *y*_*ij*_. This regression task is usually modeled as point prediction, i.e., the models only estimate the mean of the conditional distribution 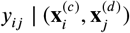 of the drug response value by optimizing the mean squared error (MSE) or a similar loss function. The conditional mean provides a narrow representation of the statistical characteristics of the response, and model certainty can only be assessed globally using empirical risk measures, such as the MSE over a test set. This assumes that the residual variance is approximately homoscedastic over the feature space. In practice, the predictability of a drug-cell line pair’s response can vary significantly depending on the cell line’s molecular state and its interactions with drug characteristics. We consider models that capture higher-order characteristics such as the variance or quantiles of the conditional distribution by explicitly accounting for heteroscedastic prediction errors. These characteristics allow the models to quantify instance-level aleatoric uncertainty in predictions.

Aleatoric uncertainty quantifies the irreducible variability in the drug response *y*_*ij*_ for a given cell line-drug pair (*i, j*) with features 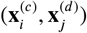, arising from inherent randomness in the system. The predicted aleatoric uncertainty is a model-derived estimate of the conditional residual variability of the response given the features. It is not a direct measurement of biological stochasticity but includes sources like measurement noise, dose–response curve-fit error, and assay artifacts. The reliability of aleatoric uncertainty breaks down in out-of-distribution scenarios. These can occur due to underrepresentation of cancer types in the training data, inconsistencies in measurement conditions, or biological shifts such as differences between patient-derived and long-cultured cell lines ***Gillet et al. (2013***). It necessitates estimating epistemic model uncertainty, which quantifies the model’s uncertainty about the predicted parameters of 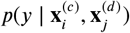, including, in probabilistic models, the predicted variance. In theory, epistemic uncertainty can be used to detect distribution shift ***Kendall and Gal (2017***). It can be reduced by collecting more data in underrepresented regions.

#### Uncertainty-Aware Models

We evaluate seven modeling approaches to assess how well different paradigms estimate uncertainty in drug response prediction. We select methods that are commonly applied, require minimal uncertainty-specific hyperparameter tuning, and produce a continuous uncertainty score. We include methods that do and do not rely on neural networks, focus on different uncertainty types (epistemic, aleatoric, or both), and span different computational costs, from linear models to neural ensembles. We exclude post-hoc wrappers on existing predictions, such as post-hoc calibrators or conformal prediction, which adjust prediction intervals but do not change the underlying uncertainty ranking ***Tibshirani et al. (2019***). Our selection covers the major families of uncertainty estimation identified in the literature ***Abdar et al. (2021); Fakour et al. (2024***): distributional modeling, ensemble methods, Bayesian inference, and evidential learning.

While in practice, the distinction between aleatoric and epistemic uncertainty is not perfect ***Gruber et al. (2025***), we categorize methods by their estimation method: distributional modeling for aleatoric, ensembles or Bayesian approaches for epistemic.

We evaluate two neural network-based approaches that estimate aleatoric uncertainty by explicitly modeling the predictive distribution:

We employ a Gaussian Neural Network (GaussNN), assuming a heteroscedastic Gaussian conditional distribution 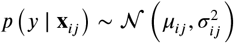, where 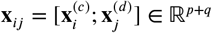 denotes the concatenated feature vector ***Nix and Weigend (1994); Bishop (1994***). We predict the distribution using a neural network *f*_*θ*_ ∶ ℝ^*p*+*q*^ → ℝ × ℝ^+^ with weights *θ* and two output neurons producing the mean and variance estimates 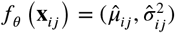. We minimize the Gaussian negative log-likelihood: 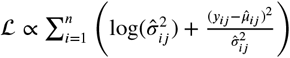 and interpret the predicted variance as a measure of the aleatoric uncertainty of the model predictions. We train it with Fast Gradient Sign adversarial training ***Goodfellow et al. (2015); Lakshminarayanan et al. (2017***) to prevent sharp predictive distributions.

Instead of modeling the output distribution parametrically, Quantile Neural Networks (QuantileNNs) predict specific quantiles, i.e., the model outputs estimates 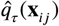 for a chosen quantile level τ ∈ (0, 1) ***Koenker and Bassett (1978); Padilla et al. (2022***). We minimize the quantile loss, which penalizes over- and underestimation of the quantile predictions. We use quantiles τ = 0.95, τ = 0.50, τ = 0.05. We measure the aleatoric uncertainty as the difference between the 0.95 and 0.05 quantile predictions (the width of the 90 % prediction interval).

To quantify epistemic uncertainty, we rely on Bayesian and ensemble-based methods. As a simple baseline, we use a Random Forest (RF). We estimate uncertainty as the variance of the predictions from individual decision trees ***Mentch and Hooker (2016***). The predictions will vary more if the models are not constrained by training examples similar to the input, and a lack of agreement, therefore, reflects epistemic uncertainty.

We also evaluate Bayesian Ridge Regression (BayesianRidge) ***MacKay (1992***), a linear model with Gaussian priors over weights and noise. It has a closed-form predictive variance Var 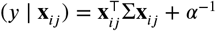, modeling epistemic uncertainty through the posterior covariance Σ and a homoscedastic component *α*^−1^.

We further employ an Monte Carlo Dropout Neural Network (MCDropout) ***Gal and Ghahramani (2016***), which applies dropout at both training and inference time. Multiple forward passes yield samples from an approximate posterior, with the variance quantifying epistemic uncertainty. We use 10 inference passes and train with MSE.

To capture both uncertainty types, we analyze a Gaussian Neural Network Ensemble (GaussNNEns) ***Lakshminarayanan et al. (2017***), where *M* = 10 GaussNNs are trained independently with different initializations. The ensemble mean serves as a point prediction, and the total predictive variance decomposes into an aleatoric component (mean of member variances) and an epistemic component (variance of member means). Mathematical details are provided in the Appendix. For this work, we also reduce inter-model correlation by resampling training and early-stopping validation data for each member.

Evidential Deep Learning (EvidentialDL) ***Amini et al. (2020***), which also uses the Gaussian likelihood, places a Normal-Inverse-Gamma prior over the likelihood parameters. The network outputs four parameters (*γ*_*ij*_, *v*_*ij*_, *α*_*ij*_, *β*_*ij*_) from which aleatoric uncertainty 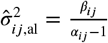 and epistemic uncertainty 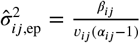 are derived, with 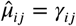 as point prediction. Training maximizes the likelihood with a regularizer that penalizes overconfidence on training examples.

#### Model Training and Evaluation

We evaluate using a 5-fold leave-cell-line-out cross-validation, to prevent cell line memorization from biasing the performance estimates. This setting reflects the primary deployment scenario of these models: predicting the response of an unprofiled cell line or patient-derived sample to a known panel of drugs. Drug response models currently still extrapolate poorly to unseen drugs (leave-drug-out setting) ***Bernett et al. (2026***). We therefore restrict our uncertainty benchmark to cell-line generalization. For hyperparameter tuning, we split an additional leave-cell-line-out validation set from the train set of each fold and perform hold-out hyperparameter optimization.

Details regarding training and hyperparameter search are in the Appendix. We assess performance of the mean estimate using the MSE and Pearson correlations between z-scaled logarithmic IC50 values and the ground truth of each cross-validation fold test set. Computing correlation metrics globally across all drugs can inflate performance due to differences in average drug potencies ***Bernett et al. (2026***). We also compute drug-wise aggregated performance metrics: We first join the test sets of all cross-validation folds, compute the metrics stratified by drug, and then report the median and standard deviation over drugs.

To evaluate uncertainties, we assess calibration and sharpness. Calibration measures how well the predicted uncertainty corresponds to the empirical test error frequency, i.e., whether prediction intervals contain the true values at the expected rate. We quantify this using the miscalibration area (MA): We integrate the absolute difference between observed and expected coverage across a grid of nominal prediction-interval levels spanning 0 to 1. The BayesianRidge, EvidentialDL, GaussNN and GaussNNEns explicitly parameterize a Gaussian predictive distribution. We approximate the outputs of other models as Gaussian, using the mean and standard deviation of predictions across ensemble members (RF, MCDropout), or by converting the interquantile range of the QuantileNN to a standard deviation. Coverage refers to the proportion of true drug response values that fall within a given prediction interval. A well-calibrated model achieves coverage close to the nominal level (e.g., a 90% prediction interval should empirically contain approximately 90% of true values). We measure sharpness by the average width of the 90 % prediction intervals. A well-calibrated model (low MA) with sharper intervals is preferred ***Chung et al. (2021***).

Calibration can be trivially improved by recalibration with a post-hoc, monotone mapping on a validation set that preserves the ordering of samples. It also does not consider how accurate the model predictions are overall (a poor model with constant uncertainties can be well calibrated). This motivates us to introduce the area under the uncertainty–reduction curve (AUURC) (smaller is better) as a metric for a joint analysis of the quality of uncertainty estimates and point predictions. The curve is constructed by ranking test instances by predicted uncertainty, and computing the MSE of the retained subset as the most uncertain instances are iteratively removed. A small area under this curve can arise from uncertainties that rank model error well or low overall point prediction errors. Conversely, larger areas indicate less informative uncertainty estimates or higher overall errors. We approximate the AUURC using numerical integration via the trapezoidal rule. We also evaluate the MSE@*k* %, i.e., the MSE when predicting only the *k* % instances with the lowest predicted uncertainty. This metric can be used to estimate the empirical risk if predictions on high-uncertainty instances are deferred. We use uncertainty-based deferral as an evaluation scenario for the informativeness of predicted uncertainties, not as a proposed decision-making strategy. In practice, uncertainty would complement domain knowledge in selecting candidates for further investigation.

We additionally report two measures of uncertainty quality that are independent of the overall point accuracy. First, we compute the Pearson correlation 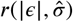 between the absolute error |*ϵ*| and the predicted uncertainty 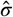. Higher correlations indicate better uncertainty quality. Second, we obtain a normalized ranking skill 1 − (AUURC − AUURC_oracle_) / (AUURC_rand_ − AUURC_oracle_). Here, AUURC_oracle_ and AUURC_rand_ are the AUURC obtained when test instances are removed in order of their absolute error (i.e., perfect removal strategy) and at random, respectively. Higher values indicate better skill of predicted uncertainty at distinguishing erroneous predictions.

For out-of-distribution detection, we add to each test feature a signed offset, with the sign drawn uniformly from {−1, +1} per feature and the magnitude drawn from a Gaussian with mean *µ* and standard deviation 10^−4^, for *µ* ∈ {0.01, 0.1, 0.2, …, 1.0}, as synthetic perturbations to the test instance features. We additionally evaluate uncertainty under real distribution shifts between screening datasets. We train each model on one screen (CTRPv2, because it relies on RNA-seq also used in the other considered study) and predict on GDSC or BeatAML. We restrict to drugs shared between the two screens and to cell lines not seen during training. We consider three transfers with different feature shifts: (i) To GDSC response data but using Broad’s CCLE RNA-seq data as features. This removes the microarray-versus-RNA-seq platform difference and leaves a biological/population shift. (ii) To the BeatAML patient extracted acute-myeloid-leukemia dataset.

BeatAML uses an independent RNA-seq screen from a different laboratory, but the same sequencing platform, introducing a cross-laboratory and biological shift. (iii) To GDSC using its native RNA microarray gene expression features. This constitutes a strong technical platform shift for the input features, in addition to the biological/population and laboratory shift. For each synthetic perturbation level and for each cross-dataset prediction, we pool the in-distribution test instances (of the CTRPv2-trained model) of all folds with the same count of matched out-of-distribution instances and use the predicted uncertainties as scores to classify each instance as out-of-distribution (high predicted uncertainty) or in-distribution (low uncertainty).

We evaluate this classification via area under the receiver operating characteristic curve (AUROC), higher values indicate stronger separation of shifted samples based on uncertainty.

### Applications of Uncertainty

#### Explaining Uncertainty

Current explainability approaches in drug response modeling attribute the predicted mean response to input features ***Kuenzi et al. (2020); Tang and Gottlieb (2021); Shi et al. (2025***). By leveraging uncertainty estimates, we can also detect drivers of predictive uncertainty. In the GaussNN, both the mean and the variance are direct outputs of the model, and we can estimate Shapley values for both outputs ***Iversen et al. (2025***). The attribution of uncertainty highlights features that act as uncertainty drivers. These features affect the models’ uncertainty estimation without necessarily influencing the predicted mean response.

#### Uncertainty-guided sample-specific fine-tuning

To evaluate whether uncertainties can be of benefit in an active learning scenario, we simulate a preclinical screening scenario in which a new cell line or patient-derived sample is added to a study. Under a limited measurement budget, the goal is to measure the drugs that maximize the information gain for the IC50 profile of a cell line or patient-derived model. This especially reflects constraints in organoid or PDX studies, where comprehensive drug screening is expensive ***Nikeghbal et al. (2025***). We pretrain a GaussNN model on a split with held-out cell lines in the test set. For each held-out cell line, we simulate measurements by selecting a set of 60 drug response observations for this cell line. We choose the drugs with the highest predicted uncertainty for that cell line as estimated by the pretrained model. As a baseline, we sample the drugs randomly from the available set. These additional drug response measurements are then combined with the original training data, and the model is fine-tuned on one of these datasets with early stopping and up to 100 epochs with the Adam optimizer defaults. We evaluate the model performance on the full set of drug responses, including the measured ones, for the held-out cell line using the MSE to measure the information gain compared to the baseline.

Further, we evaluate on the subset of drugs not included in the fine-tuning process of either method to assess the relative generalization performance increase of the baseline versus the uncertainty-guided selection.

## Results

### Benchmark

We evaluate seven models estimating drug response and aleatoric (QuantileNN, GaussNN), epistemic (RF, BayesianRidge, MCDropout), or both uncertainty types (GaussNNEns, EvidentialDL). Experiments use 5-fold leave-cell-line-out cross-validation on GDSC (924 cell lines, 367 drugs, 292,165 pairs).

#### Predictive and uncertainty performance is highest for GaussNN variants

We evaluate all models in two dimensions: predictive accuracy on unseen cell lines and the quality of their uncertainty estimates. For predictive accuracy, Table 1 reports the MSE and Pearson correlation between point predictions and ground truth. Metrics are reported either drug-wise (after merging folds, computed separately for each drug and then aggregated with the median) or globally (computed over all drug-cell-line pairs of each fold and then aggregated with the median).

**Table 1.** MSE and Pearson correlation between model predictions and experimental logarithmic IC50 values on GDSC data. For the drug-wise metrics, MSE is the median over drugs; the standard deviation represents the variability across drugs. The global metrics are calculated for the folds, and the standard deviation is the variation over folds. Best values are shown in bold; second-best are underlined. For results on CTRPv2 see Table A1.

|  | Random Forest | Bayesian Ridge | Quantile NN | MC Dropout NN | Gaussian NN | Gaussian NN Ens. | Evidential DL |
| --- | --- | --- | --- | --- | --- | --- | --- |
| *Drug-wise* |  |  |  |  |  |  |  |
| MSE | **0.187** ± 0.128 | 0.446 ± 0.596 | 0.221 ± 0.145 | 0.222 ± 0.141 | 0.202 ± 0.138 | <u>0.197</u> ± 0.141 | 0.222 ± 0.150 |
| Pearson | <u>0.425</u> ± 0.121 | 0.323 ± 0.127 | 0.331 ± 0.123 | 0.331 ± 0.122 | 0.382 ± 0.128 | **0.435** ± 0.127 | 0.325 ± 0.121 |
| *Global* |  |  |  |  |  |  |  |
| MSE | **0.220** ± 0.005 | 0.613 ± 0.005 | 0.250 ± 0.011 | 0.255 ± 0.003 | 0.237 ± 0.006 | <u>0.235</u> ± 0.004 | 0.255 ± 0.006 |
| Pearson | **0.883** ± 0.002 | 0.624 ± 0.004 | 0.867 ± 0.006 | 0.864 ± 0.002 | 0.876 ± 0.003 | <u>0.881</u> ± 0.002 | 0.864 ± 0.003 |

The reported standard deviations reflect variability across drugs or folds. The GaussNN variants and RF significantly outperform all other methods for global Pearson and MSE (paired two-sided t-tests, Benjamini-Hochberg corrected, for details see Figure A2). Bayesian Ridge consistently underperforms across all metrics, likely because of its linear model assumption. Overall, the Pearson correlations drop substantially when evaluated at the per-drug level. The large variation in overall drug potencies inflates correlations in the global setting ***Bernett et al. (2026***).

We evaluate the informativeness of the uncertainty estimates by how well the model trades off coverage of the input space suitable for predictions with the model against accuracy. Figure 2 (a) shows uncertainty reduction curves, also referred to as confidence curves ***Rasmussen et al. (2023***), for all models. Each point corresponds to the MSE obtained after iteratively removing the cell-line drug pairs in the test set with the highest uncertainty as predicted by the model and re-evaluating the MSE on the remaining samples. For informative uncertainty estimates, the curve falls monotonically (low AUURC) and achieves low MSE@*k* % at small values of *k*. By contrast, an uninformative estimate yields a near-flat trajectory, as also expected from a random order of test points. We find that the uncertainty of the BayesianRidge is uninformative, as the curve is nearly flat (see Figure A3). All other tested models exhibit a decline, indicating that higher predicted uncertainty corresponds to higher error. The GaussNN variants achieve the largest reduction. Restricting predictions to the 10 % most confident samples reduces the global MSE by 64 % (from 0.23 to 0.08) for the GaussNNEns. This reduction is not simply driven by excluding predictions far from the population mean (Figure A4). The predictions of the GaussNNEns are depicted in Figure 2 (c) and (d), with and without restricting by uncertainty to below the median.

**Figure 2.**
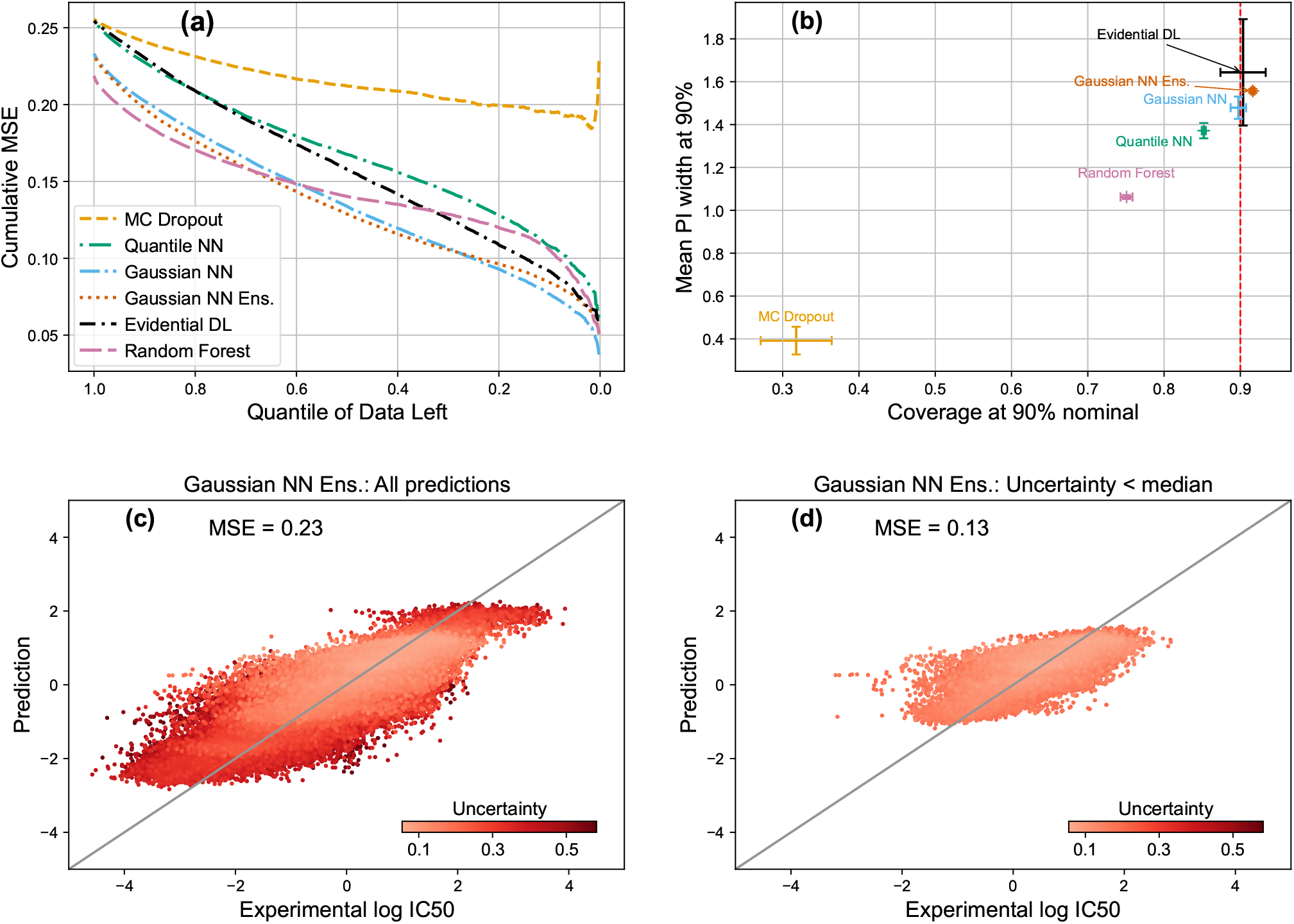
(a) Uncertainty-based filtering curves. The x-axis shows the fraction of test samples retained after thresholding by predicted uncertainty (most uncertain removed first, left to right). The y-axis reports the MSE computed on the reduced test subset. A steeper decline indicates a more effective uncertainty estimation. (BayesianRidge left out for visibility, see Figure A3) (b) Calibration–sharpness tradeoff across models. Each point is the mean empirical coverage and average prediction interval (PI) width at 90 % nominal coverage. Error bars indicate variability across folds. (c) Observed versus predicted log IC50 values for the GaussNNEns on all test samples, colored by predicted uncertainty. Same as (c), but restricted to the 50 % most confident predictions.

Table 2 summarizes the uncertainty ranking metrics AUURC and MSE@*k* % across models. The GaussNN variants achieve similar results and significantly outperform all other model types, attaining the lowest AUURC, MSE@10 %, and MSE@50 % (see Figure A5 for statistical test results). While the RF is a comparatively strong point predictor, on GDSC it trails the GaussNN variants on AUURC and MSE@*k* %. Overall, the GaussNN variants achieve the lowest MSE@10 %. The RF, EvidentialDL, and QuantileNN clearly outperform the MCDropout and BayesianRidge. For example, the MCDropout and the QuantileNN start from a similar baseline MSE (QuantileNN 0.250, MCDropout 0.255; Figure 2 (a)). Owing to better uncertainty estimation, on the 10 % most confident predictions the QuantileNN outperforms the MCDropout by 44 % (0.108 versus 0.192). The ranking skill confirms this across models: the RF and GaussNN variants are similarly accurate point predictors (Table 1), yet the RF reaches only 0.42 ranking skill, whereas the GaussNN variants rank errors best (0.51–0.52). The per-instance correlations between absolute error and uncertainty, 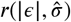, are consistent with this, delivering highest values for GaussNN variants. Note that these are modest by construction: even with perfect uncertainty estimation, the model error is a single realization drawn from the predictive distribution.

**Table 2.**
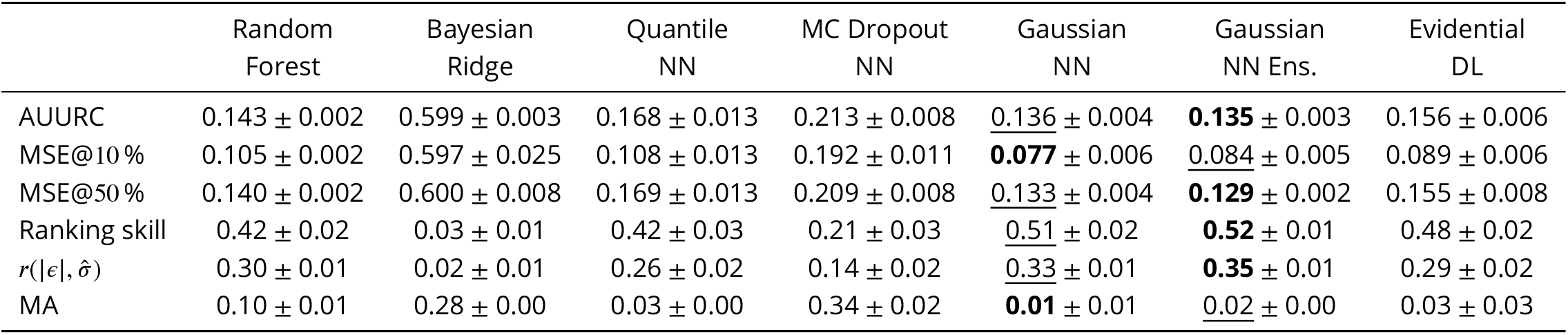
Uncertainty evaluation metrics per model on GDSC data (average ± standard deviation over the cross-validation folds). AUURC: area under the uncertainty–reduction curve (lower = better). MSE@*k* %: MSE on the *k* % most confident predictions (lower = better). MA: miscalibration area (lower = better calibrated). Ranking skill: normalized risk–reduction skill, independent of overall accuracy (1 = oracle error ordering, 0 = random; higher = better). 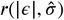 Pearson correlation between absolute error and predicted uncertainty (higher = better). Best values are shown in bold, second-best underlined. For a results on CTRPv2 data, see Table A2.

Figure 2 (b) compares the empirical coverage as a measure of uncertainty calibration and mean width of 90 % prediction intervals as a measure of sharpness of uncertainty estimates across models. Coverage close to a chosen target level (we depict the target level 90 %) with narrow prediction intervals indicates good calibration and sharpness. The GaussNN and its ensemble, as well as EvidentialDL achieve coverage near the nominal target. The calibration of these models is stable over different levels, as they achieve a low MA (see Figure A6). MCDropout and RF yield the sharpest intervals; however, only at the cost of overconfidence: Their prediction intervals contain a smaller proportion of true values than intended.

These results also largely hold on CTRPv2 (Tables A1 and A2): the GaussNN Ensemble again ranks errors best (highest ranking skill and 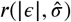). However, the RF is the strongest point predictor on CTRPv2 and, in contrast to GDSC, its tree-variance uncertainty is well calibrated for in-distribution prediction (MA ≈ 0.01).

#### Ensemble agreement is key for out-of-distribution detection

We use the AUROC to quantify how well uncertainty scores distinguish out-of-distribution (OOD) from in-distribution samples (higher values indicate better separation). As seen in Figure 3 (a) and Figure A7, the epistemic component of the GaussNNEns, and BayesianRidge are the most responsive to small synthetic perturbations and show an increase for larger shifts. MCDropout shows a similar but smaller increase. In contrast, RF tree variance remains mostly constant until a sudden rise at shift levels beyond 0.4, potentially due to the fixed thresholding in the decision trees. Aleatoric uncertainty estimates (e.g., QuantileNN, GaussNN) are considerably less affected by the shifts. To analyze more realistic OOD-scenarios, we use a CTRPv2-trained model with RNA-seq-based gene expression features on other screens. The detection AUROC rises with the severity of the real shift (Figure 3 (b)). The strongest shift is the RNA-seq-microarray platform difference, which is detected by all models except for the GaussNN. BeatAML also natively uses RNA-seq, so the shift is less severe and represents laboratory differences and a biological shift to patient-extracted leukemia samples, and is flagged by all models. Harmonizing CTRPv2 and GDSC drug response data onto shared RNA-seq gene-expression features leaves only biological/population differences, which is weakly flagged (AUROC ≈ 0.6). The prediction errors increase along with the uncertainty (Table A3). Apart from the single GaussNN, the predicted uncertainties respond to cross-dataset shifts and scale with the shift severity.

**Figure 3.**
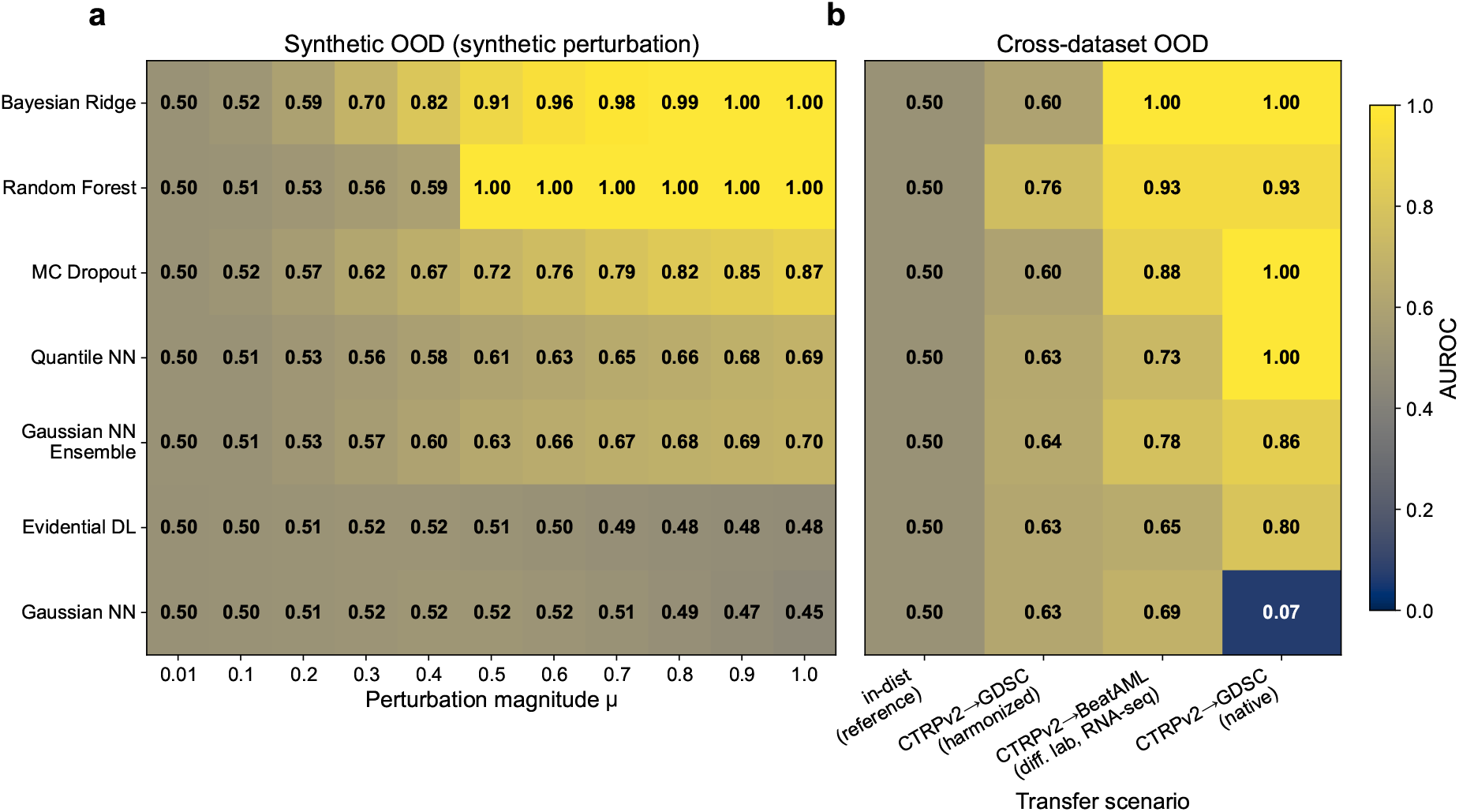
Out-of-distribution detection: AUROC for distinguishing infrom out-of-distribution samples based on predicted uncertainty. (a) Synthetic shift: perturbations of increasing magnitude added to the GDSC test features. (b) Real cross-dataset shifts of increasing severity for a CTRPv2-trained model on CCLE RNA-seq gene expression features that predicts cross-dataset with leave cell-line out split. First column: in-distribution reference. Second column: prediction on GDSC data using Broad’s CCLE RNA-seq expression features for harmonizing both datasets (biological/population difference only). Third column: on BeatAML, a different-laboratory RNA-seq cohort (same sequencing platform but different lab, and leukemia patient-extracted vs. cancer cell line samples). Fourth column: on GDSC samples with their native RNA microarray gene expression features (strong technical platform shift with biological/population difference). Model errors also rise with the increasing shift (Table A3). Epistemic uncertainties (for example, RF, BayesianRidge, MCDropout) are more reliable OOD detectors than aleatoric estimators. For standard deviations of the synthetic perturbations, see Supplementary Figure A8.

Among the tested models, the GaussNNEns best combines high predictive accuracy with calibrated aleatoric uncertainty and an epistemic component that is responsive to distributional change. However, training and inference times scale linearly with the ensemble size, making the GaussNNEns an order of magnitude slower to train than the single GaussNN (for computation time comparisons, see Table A4).

### Applications

Uncertainty-aware models enable probabilistic statements about individual predictions that go beyond point estimates. For example, rather than comparing drugs by their predicted mean response alone, one can evaluate candidates by their probability of exceeding a relevant sensitivity threshold (Figure 1 and Figure A1). We examine further uses of uncertainty estimates below.

#### Uncertainty varies systematically across tissues and drug target pathways

We investigate the potential for uncertainty estimates to provide biological insights and inform experimental design. Our analysis focuses on the GaussNN variants, which achieved the lowest overall error and most informative uncertainty estimates, trained on GDSC data. First, we examine whether predictive uncertainty varies systematically across biological and pharmacological subgroups. Figure 4 shows the distribution of the predictive uncertainty for the GaussNNEns stratified by tissue type (a) and drug target pathway (b). We observe significant differences between some tissue types (Kruskal–Wallis to test for overall differences, followed by Dunn’s test with Holm correction for pairwise comparisons). For example, the predictions in cell lines derived from hematopoietic and lymphoid tissue tumors, lymphoma and leukemia, exhibit the highest uncertainties on average, suggesting that these tissue types are more challenging to model. Notably, they are not underrepresented in the training data (Figure 4 bottom). Instead, their high uncertainty may reflect aleatoric noise or the high intra-class heterogeneity ***O’Connor and Tobinai (2014***). We observe a similar pattern for the drug-target pathway stratification. The highest median uncertainty is predicted for drugs that target the well-represented class of DNA replication (e.g., Gemcitabine, Bleomycin). In contrast, hormone-related drugs (Bicalutamide, Tamoxifen, AZD3514) are underrepresented but associated with lower predictive uncertainty, possibly due to clearer molecular determinants of response (e.g., estrogen receptor, ESR1, or androgen receptor, AR, expression levels). We analyze whether these patterns are artifacts of the underlying dose-response assay. The uncertainty does correlate with some assay quality metrics such as curve fit *R*^2^; however, this effect largely disappears once the response is accounted for, as the assay metrics are proxies for the response (Figure A10).

**Figure 4.**
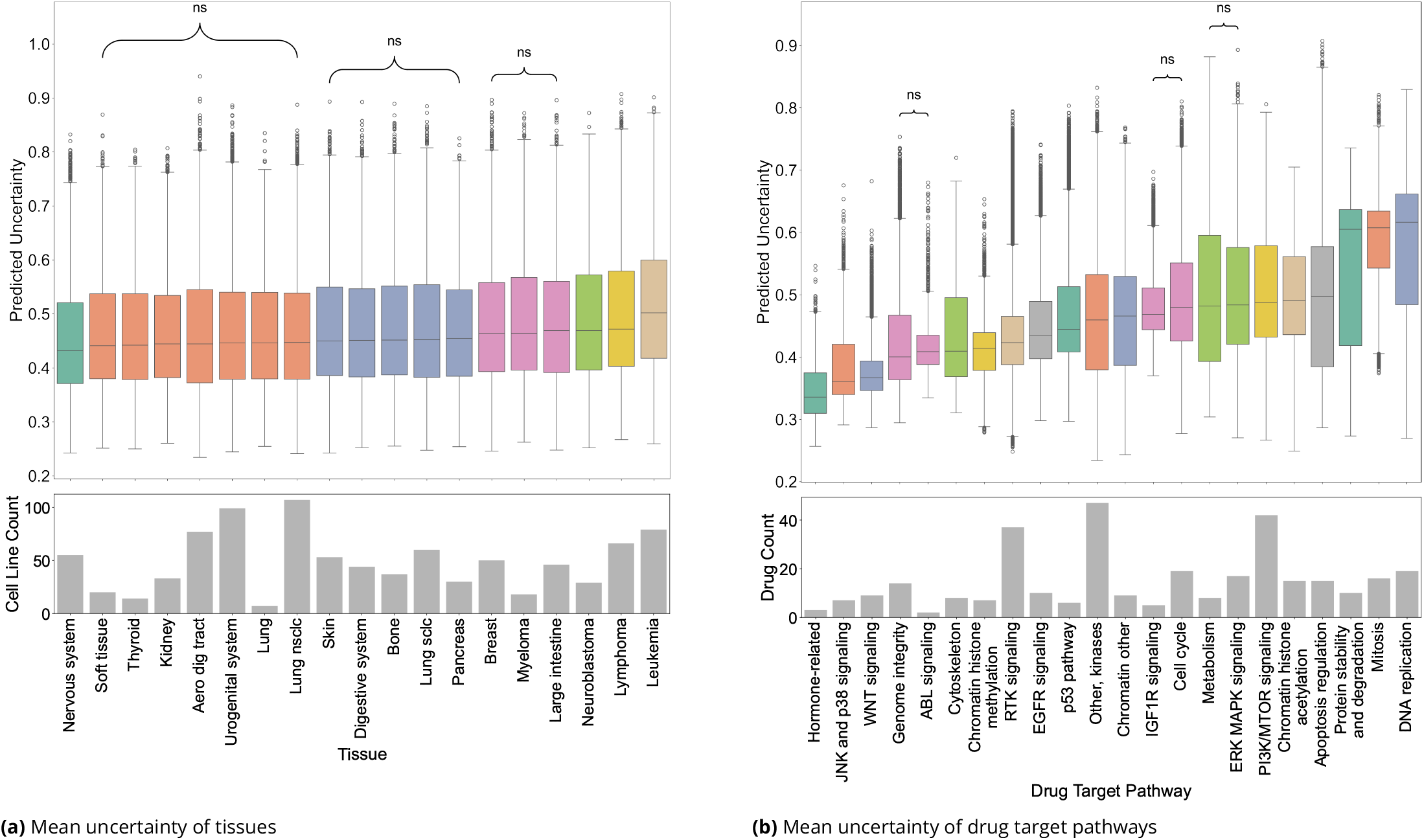
Comparison of average predictive uncertainty across (a) tissues and (b) drug target pathways of the GaussNNEns on GDSC data. Statistical differences between groups were assessed using post hoc pairwise Dunn’s test with Holm correction. Groups with non-significant pairwise differences are marked with ‘ns’ and the same boxplot color. Lower panels show sample counts per group. Predictions for hematopoietic malignancies and drugs targeting DNA replication exhibit the highest uncertainty mean. For a version of this figure with decomposed uncertainties (epistemic, aleatoric) see Figure A9.

Further, we form a tissue by pathway matrix of mean predictive uncertainty and apply hierarchical clustering (Figure 5). This analysis shows similarities between tissues and pathways, and notable outliers. Drugs affecting the ERK-MAPK signaling pathway (e.g., Selumetinib, Refametinib) are associated with high uncertainty for skin cancers. A high prevalence of ERK–MAPK pathway alterations in melanomas, particularly BRAF mutations, is known to give rise to diverse molecular subtypes with heterogeneous responses to drugs targeting ERK-MAPK ***Corrales et al. (2022***).

**Figure 5.**
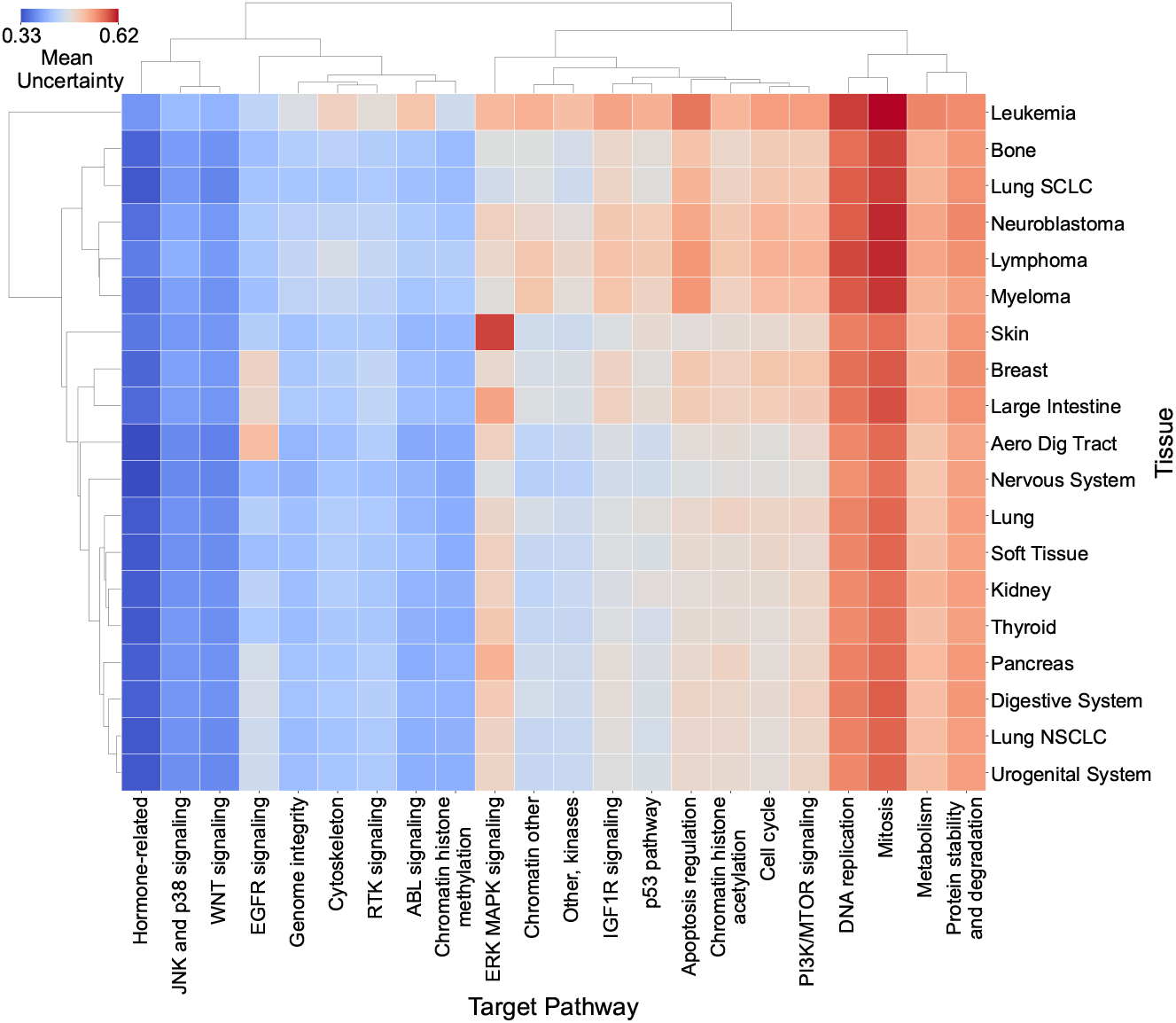
Mean predictive uncertainty across tissues and target pathways of the GaussNNEns on GDSC data, hierarchically clustered in both dimensions. Certain tissue–pathway pairs, such as ERK–MAPK signaling drugs on skin cell lines, show elevated uncertainty relative to other pairs involving these entities.

#### Uncertainty attribution reveals transcriptomic drivers of uncertainty

In Figure 6, we show the averaged mean and variance attributions of the transcriptomic input over all test instances of one cross-validation fold for the GaussNN on GDSC data. Here, genes with positive attribution on the mean output, such as the extracellular regulators of calcium-dependent signaling CALCA, MFAP5, and NMUR2, drive predictions towards resistance. In contrast, sensitivity is associated with FAM26F, GPR82, and FAM19A2, which are also part of the extracellular communication network. Genes with positive attributions for the uncertainty output, for example, signaling genes TRPM5, MESP2, and GNG2, act as drivers of unpredictability. Conversely, the genes PSMA8, TNFSF11, and GABRR1 that regulate tissue-specific functions have strong negative attributions and can be interpreted as stabilizing features. Notably, TRPM5, a transient receptor potential channel linked to chemotherapy resistance and poor prognosis in some cancers, only marginally affects the mean drug response prediction but strongly influences uncertainty estimation ***Maeda et al. (2017); Hu et al. (2023***). This modulation of uncertainty would not be captured in analyses restricted to mean effects, and provides a basis for mechanistic hypotheses about unpredictability of drug response (see also Figure A11 for a comparison of the top contributors to mean and uncertainty outputs). This attribution approach serves as a general framework that can be applied in a highly targeted manner to any tissue–pathway combination of interest. As an example, we contrast leukemia and nervous system cell lines treated with ERK-MAPK-targeting drugs (Figure A12). This analysis yields distinct gene-level drivers of both mean response and uncertainty. It allows for the generation of context-specific mechanistic hypotheses.

**Figure 6.**
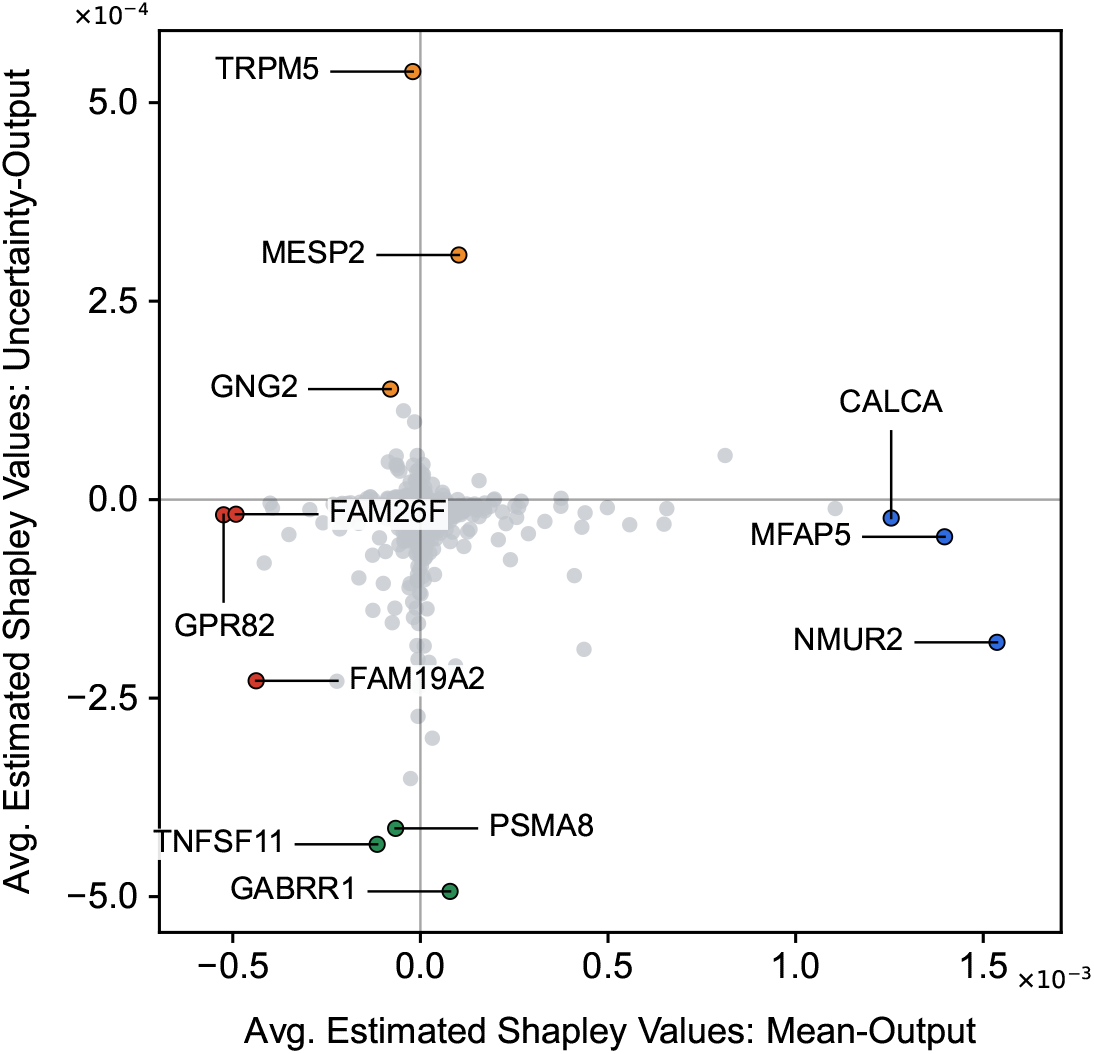
Estimated Shapley values of gene-expression features for predicted mean drug response and predictive uncertainty for the GaussNN on GDSC data averaged across the test set. The strongest average contributions, driving weak response, strong response, uncertainty, and certainty are marked in blue, red, orange, and green, respectively. Notably, some genes like TRPM5 and GABRR1 barely influence the mean predicted drug response but strongly affect the uncertainty estimate.

#### Uncertainty can inform active selection of informative measurements

We assess whether predictive uncertainty can guide the sample-specific acquisition of informative measurements to fine-tune drug response models. This setting reflects scenarios in which drug response measurements are costly, as in organoid or PDX experiments, and only a limited number of compounds can be screened for a new sample. On the full drug set, including measured drugs, uncertainty-guided fine-tuning significantly reduces MSE compared to random selection (mean MSE 0.216 vs. 0.224, *p* < 10^−16^, rank-biserial *r* = 0.60, two-sided Wilcoxon signed-rank test, *n* = 920, see Figure A13). This result is consistent with the greedy strategy targeting the most uncertain samples, confirming that the uncertainty estimates meaningfully identify where the model has the most to gain. A small improvement persists even when evaluation is restricted to the intersection of drugs not used for fine-tuning for both approaches (mean MSE 0.183 vs. 0.184, *p* = 2 × 10^−4^, rank-biserial *r* = 0.14, *n* = 920). Research into balancing this exploitative strategy with diversity-oriented sampling could lead to greater improvements in this setting.

## Discussion

We benchmark several models that predict drug response along with their predictive uncertainty. We find that distributional modeling methods (QuantileNN, GaussNN) are effective for ranking in-distribution errors of drug-cell line pairs. However, they are unresponsive to distributional shift, and, therefore, their uncertainty estimates are unreliable in settings involving possible domain shifts. Ensemble-based approaches (MCDropout, RF) complement this: While their uncertainty estimates are considerably less precise at ranking errors, they can signal out-of-distribution inputs, indicating the boundaries of the model’s applicability. Combining distributional modeling with ensembling (e.g., GaussNNEns) can provide both advantages, albeit with a significantly higher computational cost. It is the only calibrated model with a functional epistemic component. EvidentialDL promises a single-model alternative that captures both uncertainty types, but it achieved only moderate uncertainty-based error ranking performance and failed to detect distribution shifts, possibly due to its hyperparameter sensitivity. Taken together, our results suggest matching the uncertainty method to the application: distributional models when predictions stay within a single screening panel with consistent measurements, and ensembles when models must generalize across datasets, laboratories, or assay platforms, where batch effects and differences in experimental protocols can introduce distributional shifts. Consistent with our recent benchmark ***Bernett et al. (2026***), RF is the strongest point predictor.

Its tree-variance uncertainty, however, trails the GaussNN variants on uncertainty evaluation metrics on GDSC and detects out-of-distribution inputs only for large perturbations. This confirms that response prediction accuracy and uncertainty quality are largely orthogonal. The gap is dataset-dependent: the RF’s tree-variance uncertainty is overconfident on GDSC but well calibrated on CTRPv2. A large body of work has explored advanced modeling approaches for drug response prediction, but the evaluations focus on point prediction accuracy. We recommend assessing performance in two dimensions: accuracy and uncertainty quality (e.g., using AUURC and MSE@*k* %). For our experiments, all neural network models used simple expression- and fingerprint-based feed-forward architectures (GaussNN, GaussNNEns, QuantileNN, MCDropout). Whether more complex architectures, additional modalities, or prior knowledge can improve uncertainty quality and expose apparent aleatoric uncertainty as epistemic remains to be studied.

Modeling uncertainty is not only a measure of prediction reliability but facilitates new downstream applications. It can be used to probabilistically analyze predictions and aggregating uncertainties by tissue and pathway reveals systematic differences between tissues and drug classes. This can guide the allocation of future screening efforts: cancer types with consistently high uncertainty are recommended for expanded profiling. The ability to attribute uncertainty separately from mean predictions opens a new dimension for generating mechanistic hypotheses. Traditional explainability approaches in the drug response field identify genes associated with sensitivity or resistance, but attributing uncertainty can identify genes that drive (un)predictability. Their expression may characterize biological states that lead to variable or context-dependent drug responses. This could inform mechanistic follow-up studies to study the sources of response variability. The identified genes are distinct from standard sensitivity markers and provide orthogonal information that would be missed in analyses restricted to the mean response.

Finally, uncertainties can be used to prioritize measurements for sample-specific fine-tuning of drug response models, and lead to small but significant gains over a random selection strategy. We expect that strategies that balance exploitation based on uncertainty with exploration of the feature space could yield larger improvements.

Our analysis is subject to limitations. First, we benchmark methods on the widely used GDSC dataset. While we confirm the results for the CTRPv2 dataset, both rely on cell lines which are long-cultured *in vitro* cancer models that differ from primary, patient-extracted cell lines. In addition, we summarize drug sensitivity with the IC50, which yields no direct information about the therapeutic index. Translating uncertainty-aware drug response models into clinical decision support requires several steps beyond this preclinical benchmark. Second, uncertainty-based deferral may preferentially exclude highly sensitive cell line–drug pairs, biasing the retained subset toward easier but less actionable cases. This may also mean that aggregate performance metrics may overstate the ability to identify the most responsive pairs. Developing evaluation metrics that account for this reliability–informativeness tradeoff is an open challenge. Further, the analysis of drivers of drug response and uncertainty we conduct does not establish causality. The Shapley values here reflect contributions from the trained model, not necessarily biological effects. High attribution scores can arise via correlations with confounding factors such as tissue or cell line identity, rather than mechanistically relevant genes. Future work could identify drivers that are independent of cell line or tissue confounders, for example, by applying tissue-invariant risk minimization that controls for tissue or cell line context. The stability of the uncertainty drivers across independent datasets remains to be established.

Despite these limitations, we recommend uncertainty estimation as standard practice in drug response modeling. Our results show that uncertainty estimates enable qualitatively new analyses, from identifying molecular drivers of unpredictability to prioritizing experiments under resource constraints. As drug response models move toward more complex and increasingly multimodal architectures, systematic evaluation of uncertainty quality in addition to predictive accuracy will be important for building models that are more transparent about their limits.

## Ethics Statement

This study uses only publicly available cell line screening data from the Genomics of Drug Sensitivity in Cancer (GDSC) project and the Cancer Therapeutics Response Portal (CTRPv2). We additionally use the BeatAML dataset of ex vivo drug-sensitivity measurements on de-identified primary tumor samples from acute myeloid leukemia patients. All data were generated by the original studies under their respective ethics approvals. No ethics approval was required.

## Data Availability

The GDSC data are publicly available at https://www.cancerrxgene.org/. The CTRPv2 drug response data are available through the Cancer Therapeutics Response Portal (https://portals.broadinstitute.org/ctrp.v2/). The CCLE RNA-seq gene expression data are available from the Broad Institute CCLE (https://sites.broadinstitute.org/ccle/datasets). The BeatAML *ex vivo* drug response and RNA-seq gene expression data are available through the Beat AML data viewer (http://www.vizome.org/). The code, cross-validation splits, processed input data, and model outputs are available at https://github.com/PascalIversen/LUDRP and Zenodo (https://doi.org/10.5281/zenodo.19219090). For the cross-dataset experiments, we use the preprocessed datasets from DrEval (https://doi.org/10.5281/zenodo.12633909).

## Appendix

### Training details

If not mentioned otherwise, experiments are conducted using the Genomics of Drug Sensitivity in Cancer (GDSC) dataset. RNA microarray gene expression features are arcsinh-transformed and z-normalized using train set statistics. Drug features are represented as 128-bit Morgan fingerprints. The final input vector is obtained by concatenating gene expression and drug features. The target variable is the logarithmic z-scaled IC50.

Model training and evaluation follows a five-fold cross-validation protocol, where we exclude cell lines in the test from the training set. Within each fold, a leave-cell-line-out validation set is split from the training set for inner hold-out hyperparameter optimization with exhaustive grid search.

The models are fitted on the full train set with the hyperparameter that yielded the lowest-MSE after hyperparameter optimization. Model performance metrics are then estimated on the test set.

For neural network models (QuantileNN, MCDropout, GaussNN, GaussNNEns), we additionally split off a leave-cell-line-out early-stopping-validation set from each training set. We set a maximum of 1000 Epochs and stop the training when the early-stopping-validation loss ceases to improve for five epochs (patience parameter), and restore the best model with respect to the early-stopping-validation loss after the training stops. We implement all neural models in PyTorch, use ReLU activation, and train with the Adam optimizer with default parameters and a batch size of 128. The ensemble model (GaussNNEns) uses fixed architectures of [128, 32, 16] with dropout 0.3 and 10 ensemble members to reduce computational cost. For the CTRPv2 replication and cross-dataset, we use 5 ensemble members to reduce the computational cost. For all other neural network models, we conduct hyperparameter optimization over network architectures with hidden layer configurations of [64, 16, 8], [128, 32, 16], and [64, 16, 16, 8, 8] units, combined with dropout rates of 0.1 and 0.3. For the EvidentialDL model, we use the same architecture search space and additionally optimize the evidence regularization weight over λ ∈ {10^−2^, 10^−1^}. We verified that a wider search (λ ∈ {10^−3^, 10^−2^, 10^−1^, 1}) does not improve EvidentialDL. The GaussNN and GaussNNEns are trained with adversarial training following the Fast Gradient Sign Method (FGSM) ***Goodfellow et al. (2015); Lakshminarayanan et al. (2017***). At each training step, an adversarial example is generated by perturbing the input x as x_adv_ = x + *ϵ* ⋅ sign(∇_x_ *L*(x, *y*)), with *ϵ* = 0.01. The total training loss is the sum of the standard loss and the loss on the adversarial example. Classical machine learning models (BayesianRidge, RF) were implemented using scikit-learn. We optimize Bayesian Ridge regression over precision parameters *α*_1_ ∈ {10^−6^, 10^−5^, 10^−4^}, *α*_2_ ∈ {10^−6^, 10^−4^}, with fixed λ_1_ = λ_2_ = 10^−6^ . For the Random Forest, we explore forests of 200 and 500 trees with maximum depths of 10 and 15, each tree trained on a random 85% of the samples and 85% of the features (row and feature subsampling); the best configuration (500 trees, maximum depth 15) is selected per fold. For the CTRPv2 replication, the Random Forest uses this configuration without further tuning. The code, the cross-validation splits, and the input data for all experiments and evaluations are available at https://github.com/PascalIversen/LUDRP. Model parameter count and run times are listed in Table A4.

### Gaussian NN Ensemble variance decomposition

Let *M* denote the number of ensemble members and *m* ∈ {1, …, *M*} the member index. Each member outputs 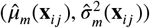. The ensemble prediction and uncertainty decomposition of the

GaussNNEns model are:

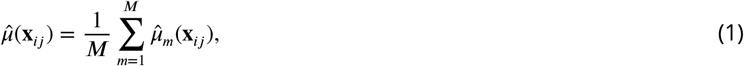

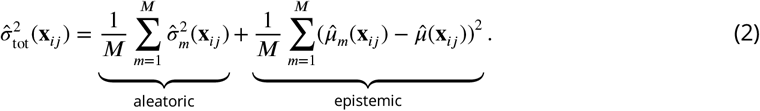

**Table A1.** MSE and Pearson correlation between model predictions and experimental logarithmic IC50 values on the CTRPv2 dataset (5-fold leave-cell-line-out). For the drug-wise metrics, MSE is the median over drugs; the standard deviation represents the variability across drugs. The global metrics are calculated for the folds, and the standard deviation is the variation over folds. Best values are shown in bold; second-best are underlined.

|  | Random Forest | Bayesian Ridge | Quantile NN | MC Dropout NN | Gaussian NN | Gaussian NN Ens. | Evidential DL |
| --- | --- | --- | --- | --- | --- | --- | --- |
| *Drug-wise* |  |  |  |  |  |  |  |
| MSE | **0.160** ± 0.487 | 0.537 ± 0.982 | 0.277 ± 0.714 | 0.279 ± 0.634 | 0.215 ± 0.575 | <u>0.198</u> ± 0.545 | 0.262 ± 1.014 |
| Pearson | **0.391** ± 0.289 | 0.261 ± 0.189 | 0.207 ± 0.213 | 0.236 ± 0.190 | 0.297 ± 0.217 | <u>0.345</u> ± 0.239 | 0.242 ± 0.205 |
| *Global* |  |  |  |  |  |  |  |
| MSE | **0.236** ± 0.008 | 0.677 ± 0.014 | 0.321 ± 0.030 | 0.330 ± 0.019 | 0.276 ± 0.008 | <u>0.274</u> ± 0.008 | 0.324 ± 0.028 |
| Pearson | **0.874** ± 0.003 | 0.571 ± 0.006 | 0.832 ± 0.014 | 0.823 ± 0.009 | 0.854 ± 0.003 | <u>0.863</u> ± 0.004 | 0.824 ± 0.015 |

**Table A2.**
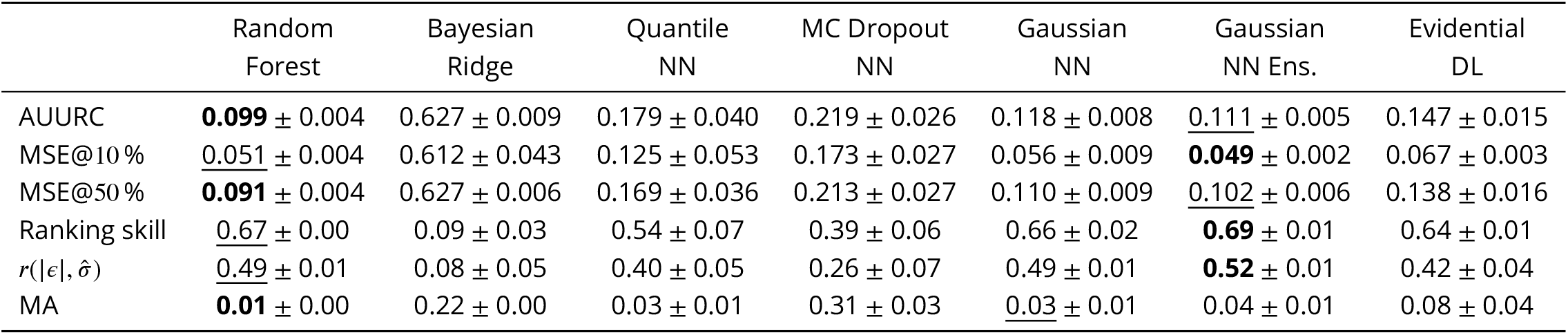
Uncertainty evaluation metrics per model on the CTRPv2 dataset (average ± standard deviation over the cross-validation folds). AUURC: area under the uncertainty–reduction curve (lower = better). MSE@*k* %: MSE on the *k* % most confident predictions (lower = better). MA: miscalibration area (lower = better calibrated). Ranking skill: normalized risk–reduction skill, independent of overall accuracy (1 = oracle error ordering, 0 = random; higher = better). 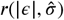 Pearson correlation between absolute error and predicted uncertainty (higher = better). Best values are shown in bold, second-best underlined.

**Table A3.**
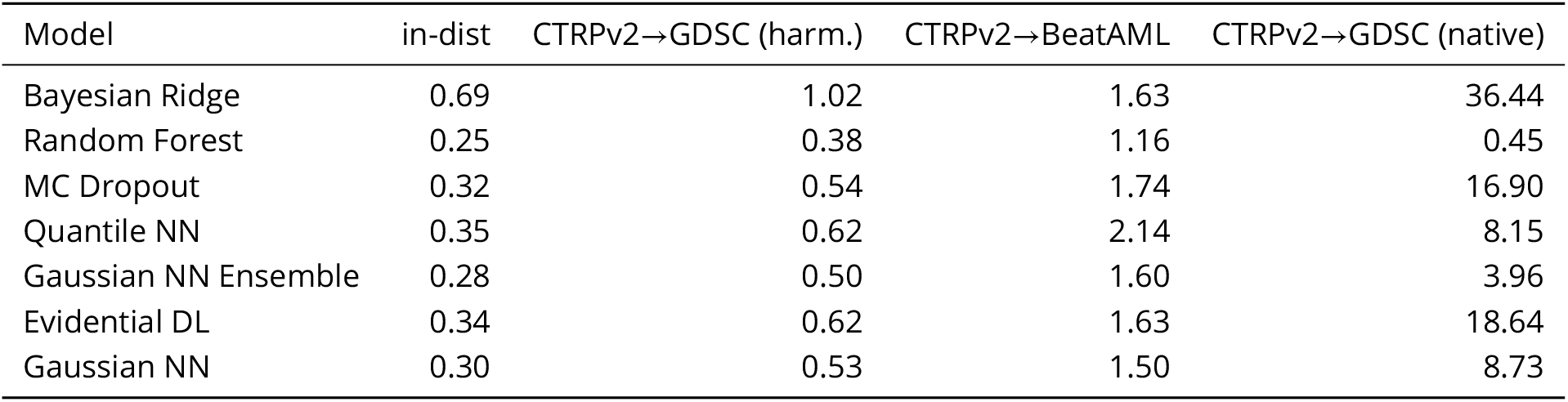
Transfer mean squared error (on the globally z-scored response, smaller is better) for the cross-dataset out-of-distribution scenarios of Figure 3(b), ordered by increasing shift. The GDSC transfers are averaged over five random cell-line splits; the BeatAML transfer uses a single split.

| Model | in-dist | CTRPv2→GDSC (harm.) | CTRPv2→BeatAML | CTRPv2→GDSC (native) |
| --- | --- | --- | --- | --- |
| Bayesian Ridge | 0.69 | 1.02 | 1.63 | 36.44 |
| Random Forest | 0.25 | 0.38 | 1.16 | 0.45 |
| MC Dropout | 0.32 | 0.54 | 1.74 | 16.90 |
| Quantile NN | 0.35 | 0.62 | 2.14 | 8.15 |
| Gaussian NN Ensemble | 0.28 | 0.50 | 1.60 | 3.96 |
| Evidential DL | 0.34 | 0.62 | 1.63 | 18.64 |
| Gaussian NN | 0.30 | 0.53 | 1.50 | 8.73 |

**Table A4.** Number of model parameters and computation time for training and test set prediction on one fold. Device: Apple MPS.

| Model | # Parameters | Train (s) | Time/Epoch (s) | Predict (s) |
| --- | --- | --- | --- | --- |
| Random Forest | — | 698.3 | — | 12.84 |
| Bayesian Ridge | — | 64.5 | — | 3.04 |
| Quantile NN | 288,611 | 94.7 | 8.61 | 0.11 |
| MC Dropout NN | 288,577 | 141.9 | 6.17 | 0.64 |
| Gaussian NN | 288,594 | 392.7 | 14.02 | 0.12 |
| Gaussian NN Ens. | 2,885,940 | 8534.3 | — | 2.06 |
| Evidential DL | 288,628 | 184.3 | 10.84 | 0.12 |

**Figure A1.**
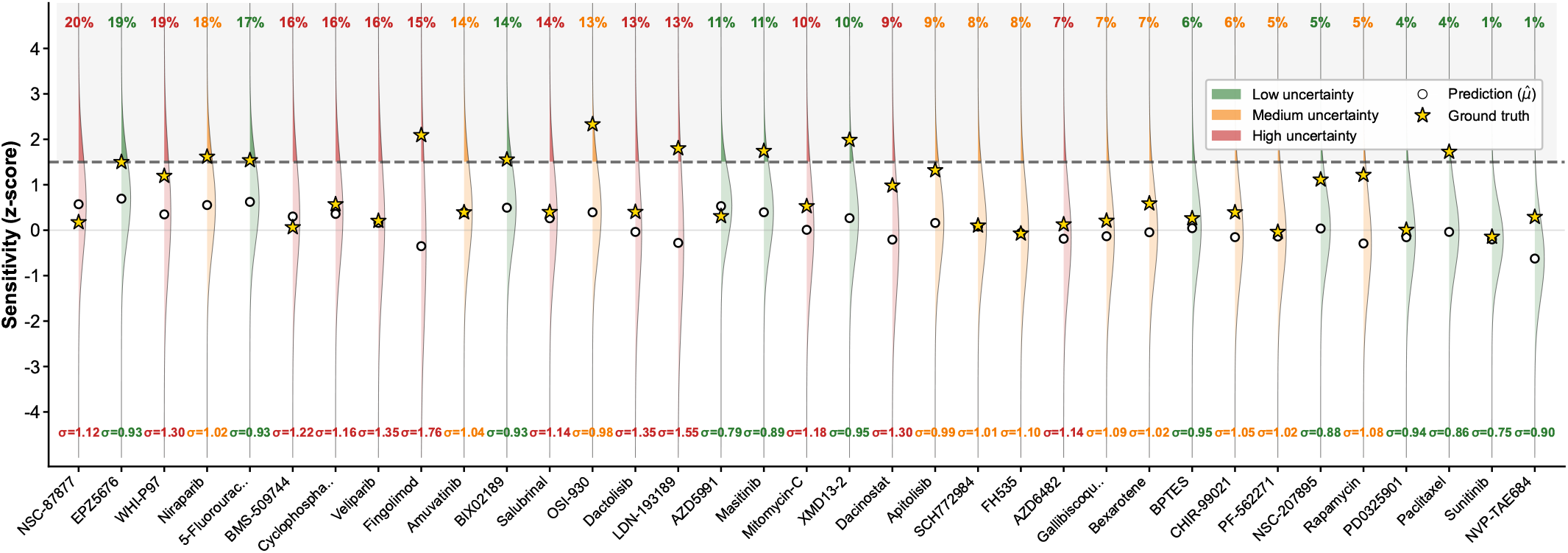
Predicted and experimental relative drug sensitivity distributions for the CESS leukemia cell line across a sample of 35 drugs. Sensitivities are the per-drug z-scored negative logarithmic IC50. Each drug’s predictive distribution of the GaussNNEns on GDSC data is shown. Drugs are sorted by the probability of exceeding a sensitivity threshold of *z* = 1.5, shown as a percentage above each drug.

**Figure A2.**
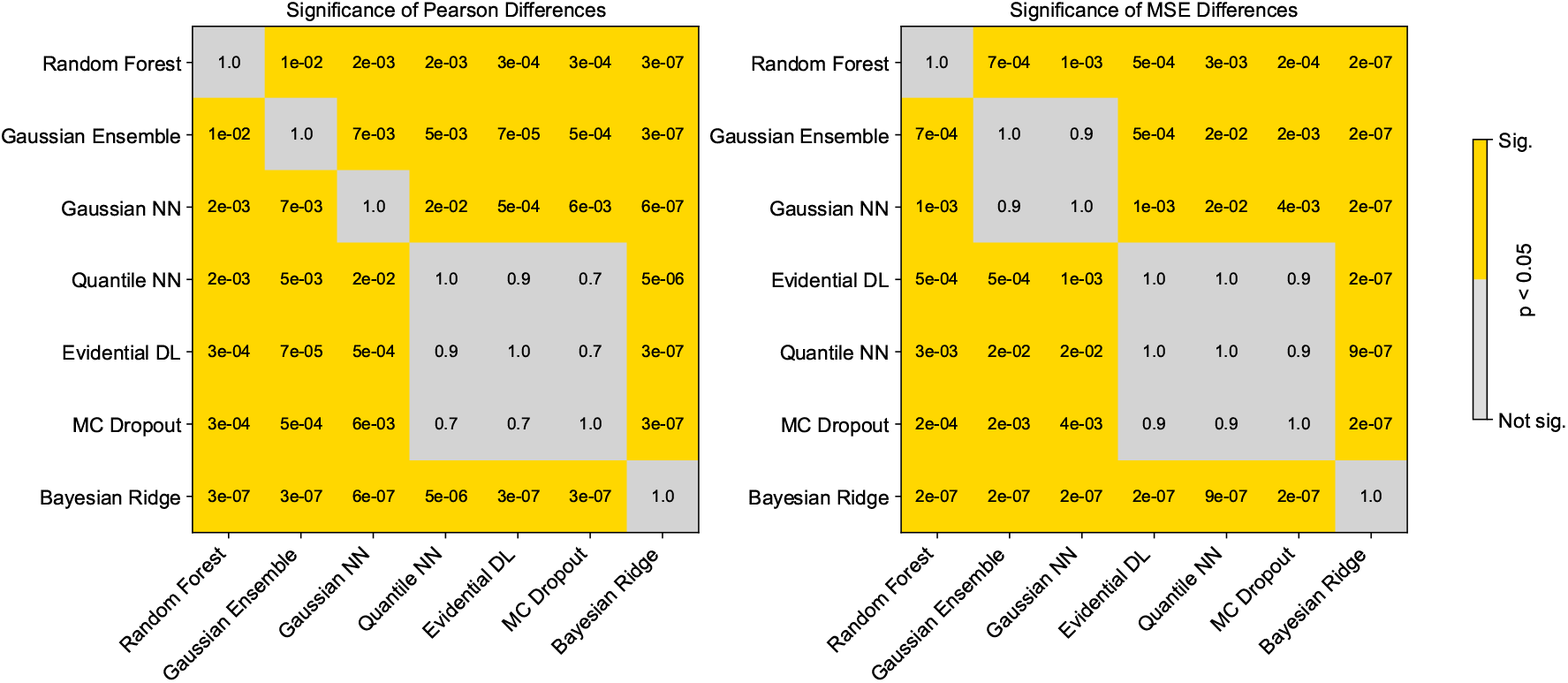
Pairwise statistical comparison of model accuracy for prediction on GDSC across metrics. Each cell shows the result of a paired two-sided t-test between two models over cross-validation folds, corrected for multiple testing using the Benjamini–Hochberg procedure. Left: significance of differences in global Pearson correlations after Fisher-z transformation. Right: significance of differences in global mean squared error. Models are sorted for performance.

**Figure A3.**
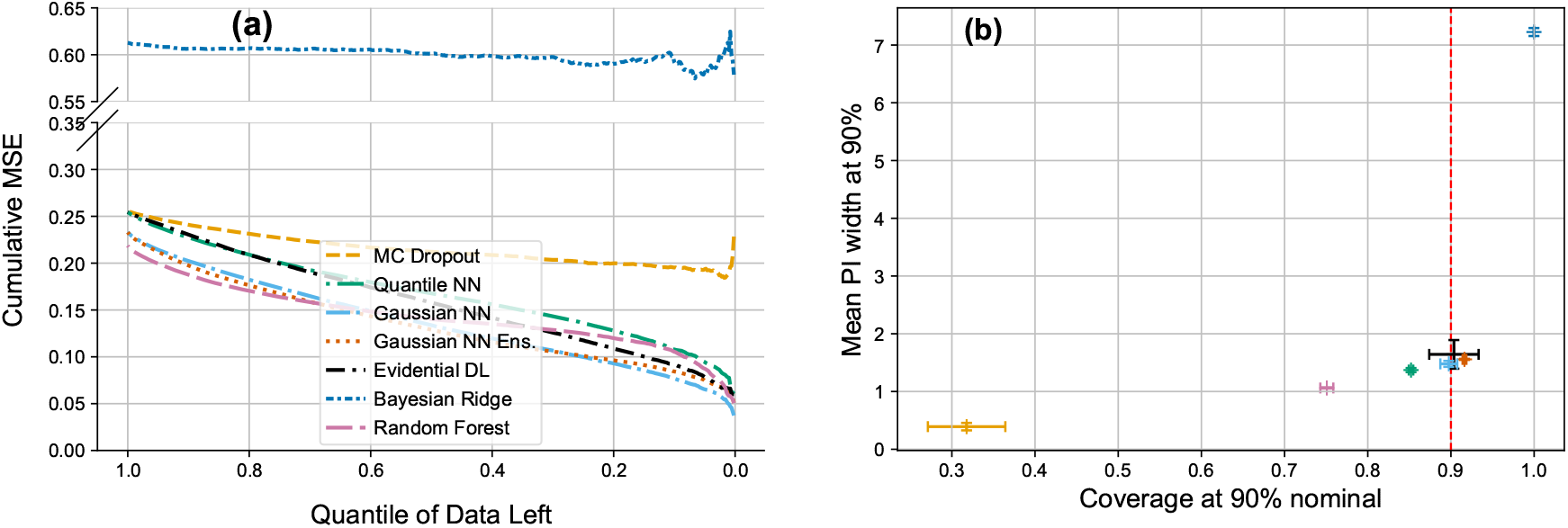
(a) Uncertainty-based filtering curves for each model on GDSC data, including Bayesian Ridge. The x-axis shows the fraction of test samples retained after thresholding by predicted uncertainty (most uncertain removed first, left to right). The y-axis reports the MSE computed on the reduced test subset. A steeper decline indicates a more effective uncertainty estimation. (b) Calibration–sharpness tradeoff across models. Each point is the mean empirical coverage and average prediction interval (PI) width at 90 % nominal coverage. Error bars indicate variability across folds.

**Figure A4.**
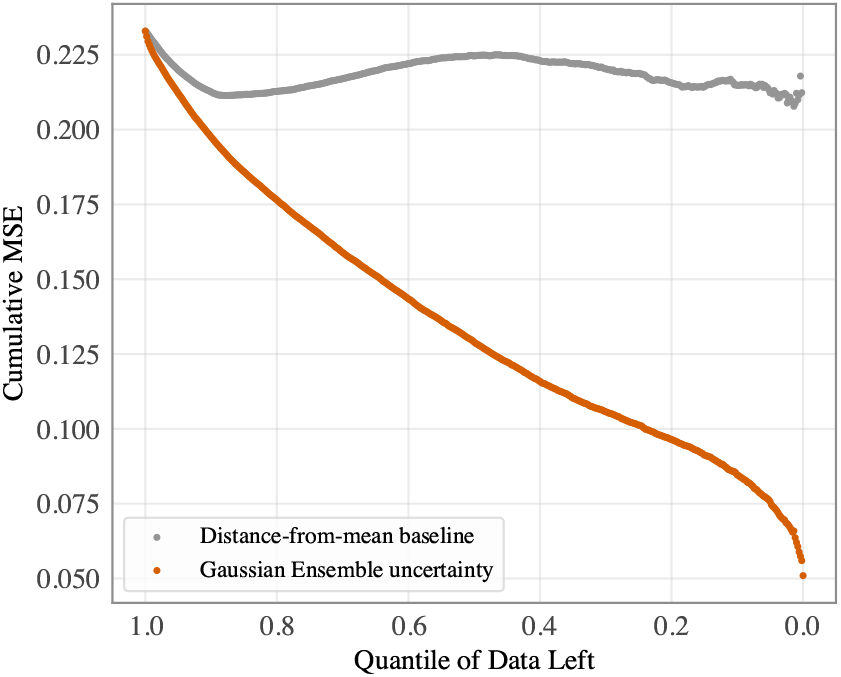
Uncertainty reduction curve for the GaussNNEns on GDSC compared to a baseline, where instances are ordered by the absolute distance of their predictions from the mean prediction.

**Figure A5.**
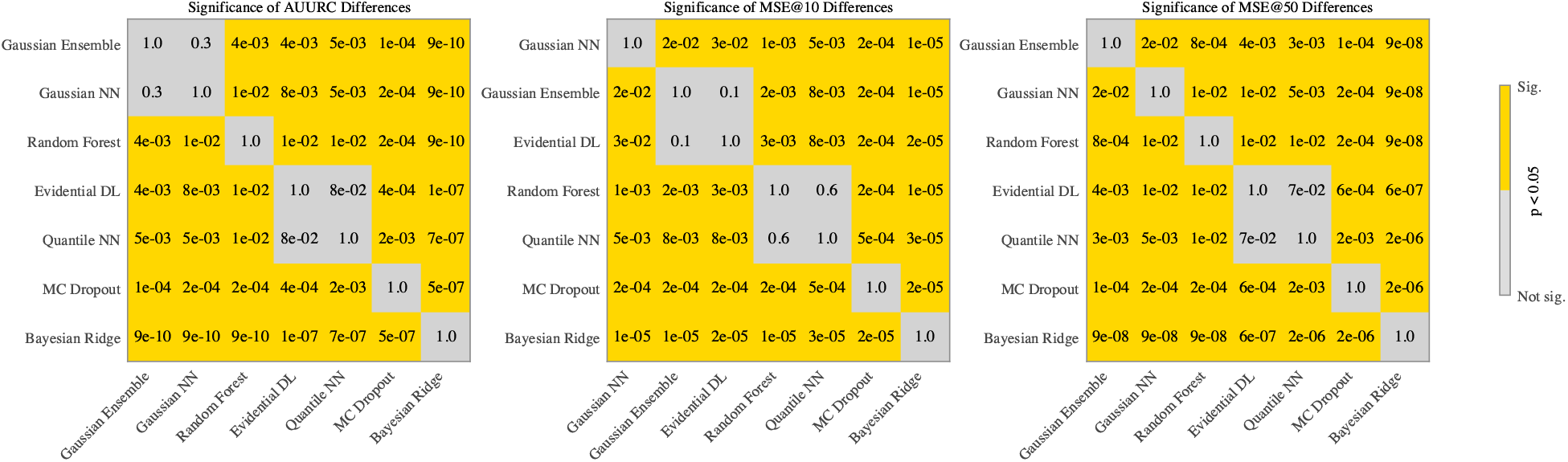
Pairwise statistical comparison of uncertainty metrics for prediction on GDSC across metrics. Each cell shows the result of a paired two-sided t-test between two models over cross-validation folds, corrected for multiple testing using the Benjamini–Hochberg procedure. Models are sorted for performance.

**Figure A6.**
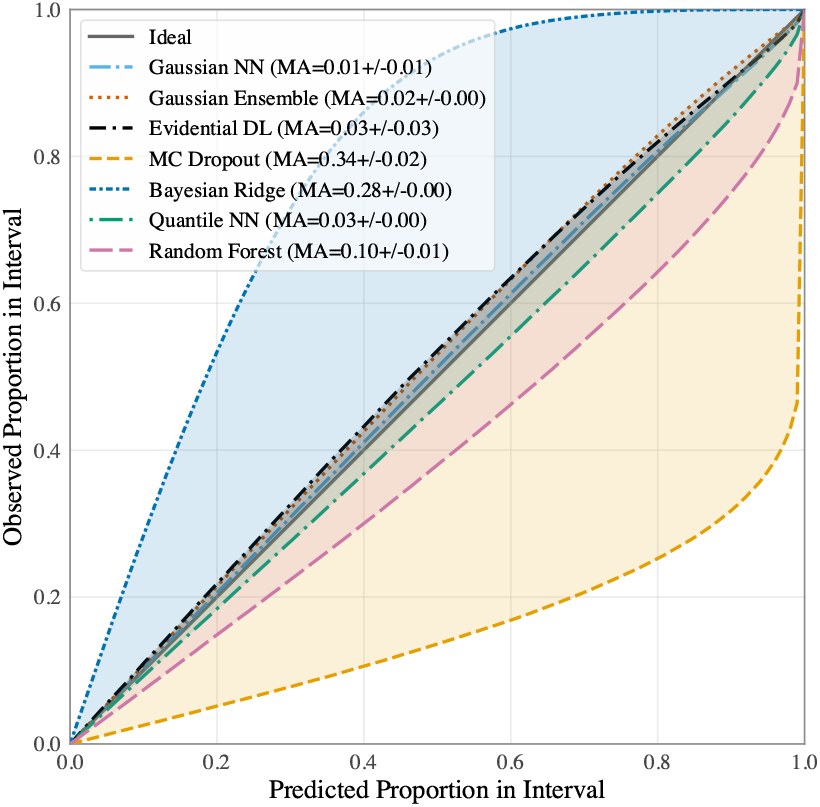
Calibration curves comparing predicted and observed proportions of responses within uncertainty intervals for each model on GDSC data. Models with curves close to the ideal diagonal are well calibrated (GaussNN, GaussNNEns, EvidentialDL, QuantileNN). Curves above the diagonal indicate underconfident predictions (BayesianRidge), while curves below indicate overconfident predictions (MCDropout, RF). Shaded regions correspond to miscalibration areas (MA) for each model. All calibration results assume a Gaussian predictive distribution.

**Figure A7.**
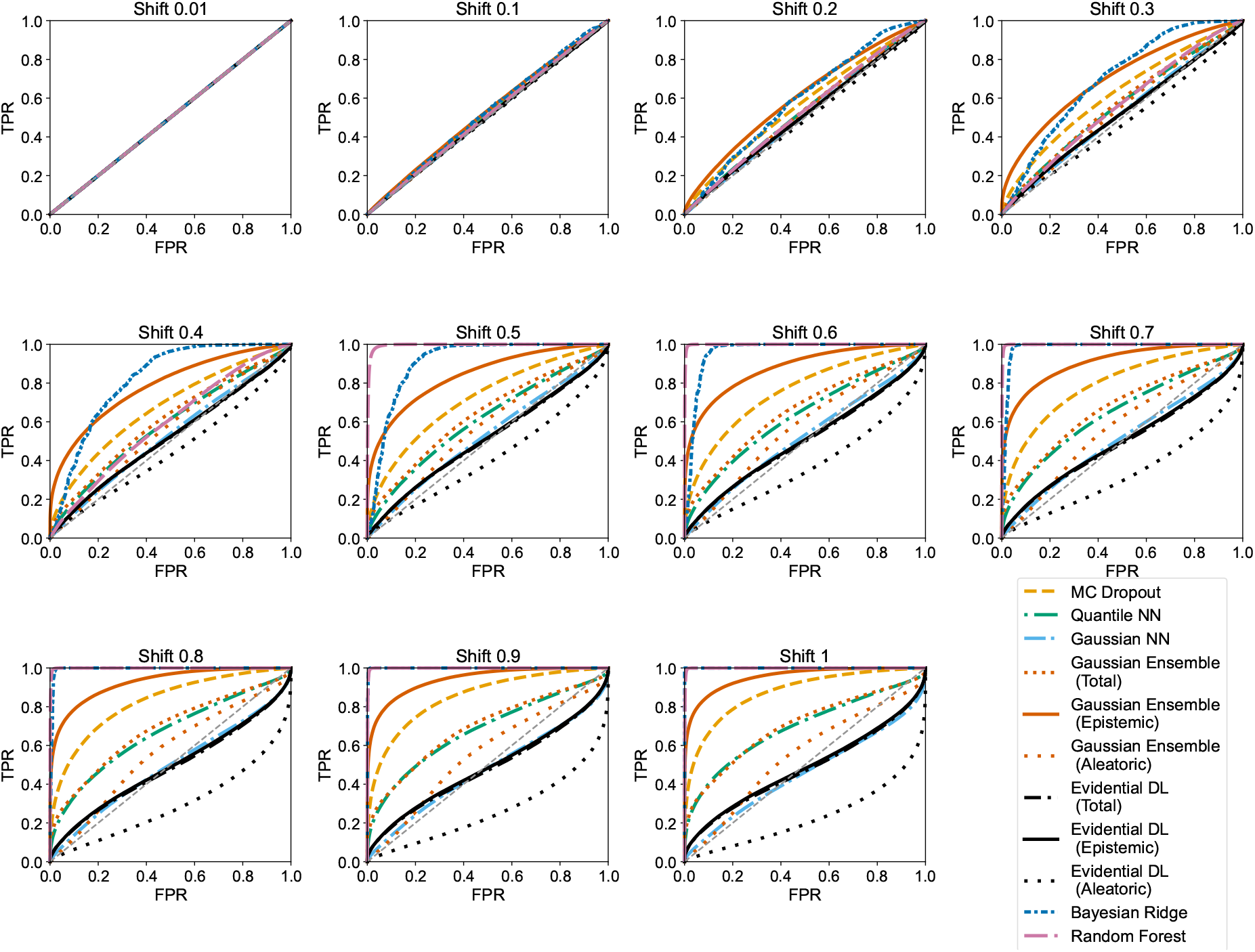
Receiver Operating Characteristic (ROC) curves for out-of-distribution detection performance on GDSC across different distribution shifts induced by synthetic perturbations. Each subplot corresponds to one shift level, showing true positive rate (TPR) versus false positive rate (FPR). We also show GaussNNEns uncertainty decomposed into epistemic and aleatoric components.

**Figure A8.**
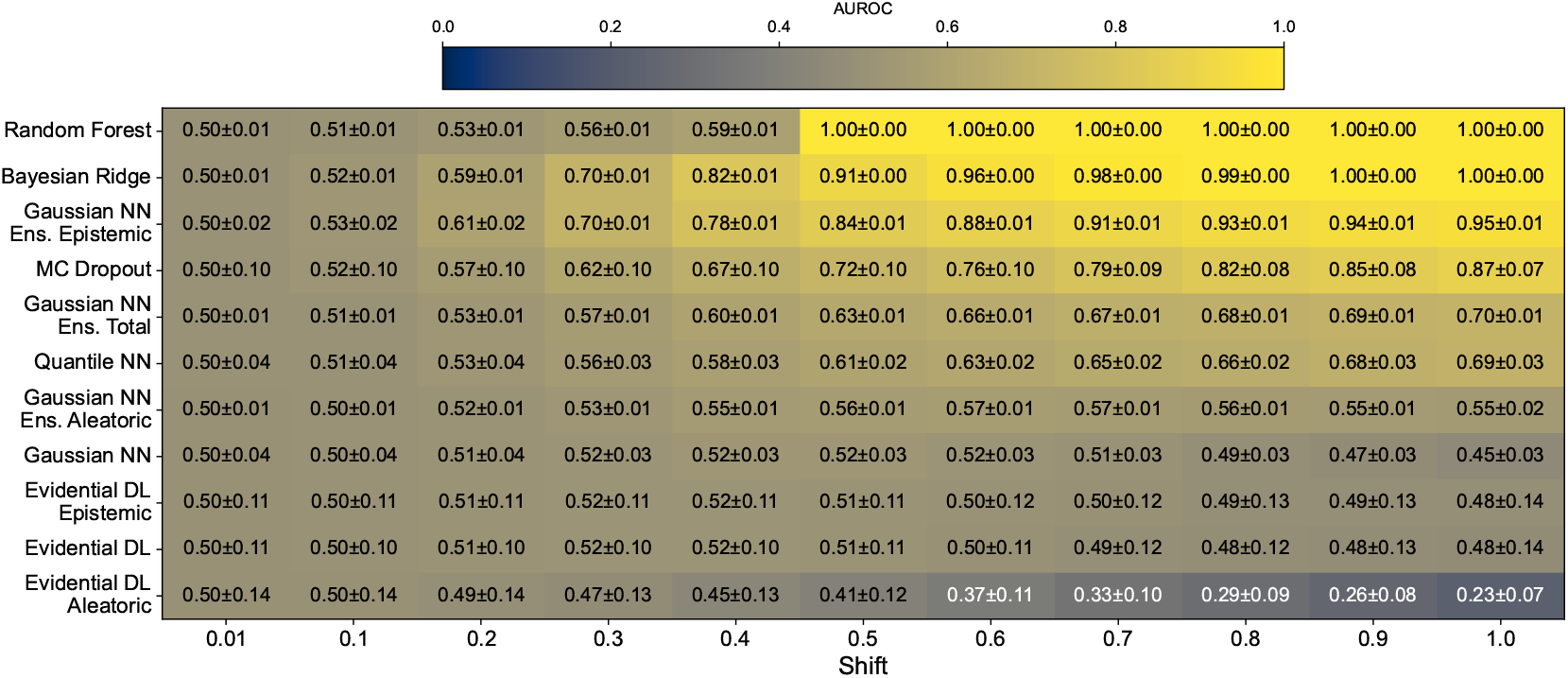
Out-of-distribution (OOD) detection on GDSC-trained models: AUROC (higher is better) for distinguishing synthetically perturbed from unperturbed samples based on predicted uncertainty for increasing distribution shift strengths. Variation is the standard deviation of the AUROC over the five cross-validation folds.

**Figure A9.**
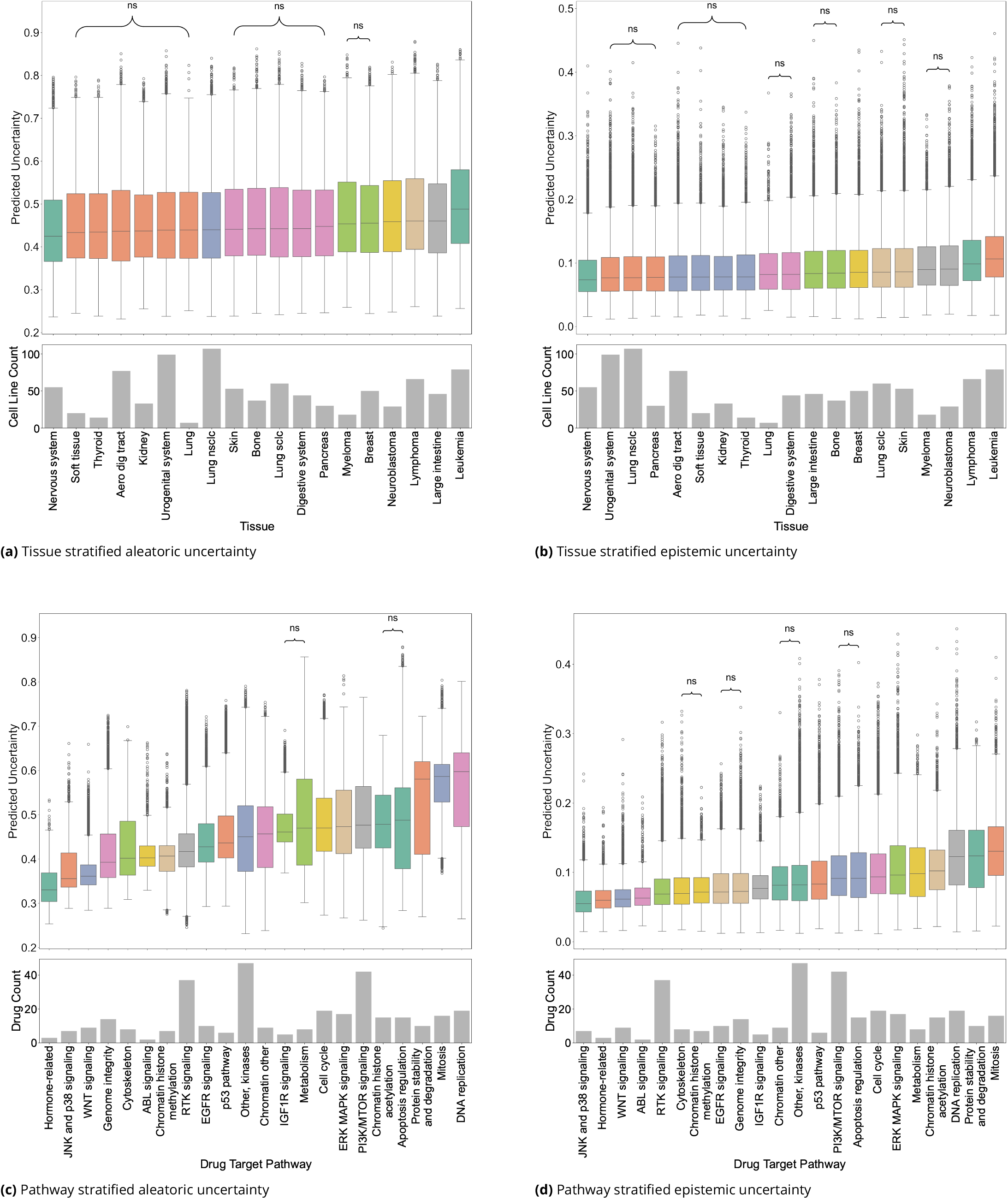
Comparison of predictive uncertainties estimated by the GaussNNEns on GDSC. (a, b) Aleatoric and epistemic uncertainty across tissues. (c, d) Aleatoric and epistemic uncertainty across drug target pathways. Statistical differences between groups were assessed using post hoc pairwise Dunn’s test with Holm correction. Non-significant pairwise differences are marked with ‘ns’. Lower panels of each subplot show sample counts per group.

**Figure A10.**
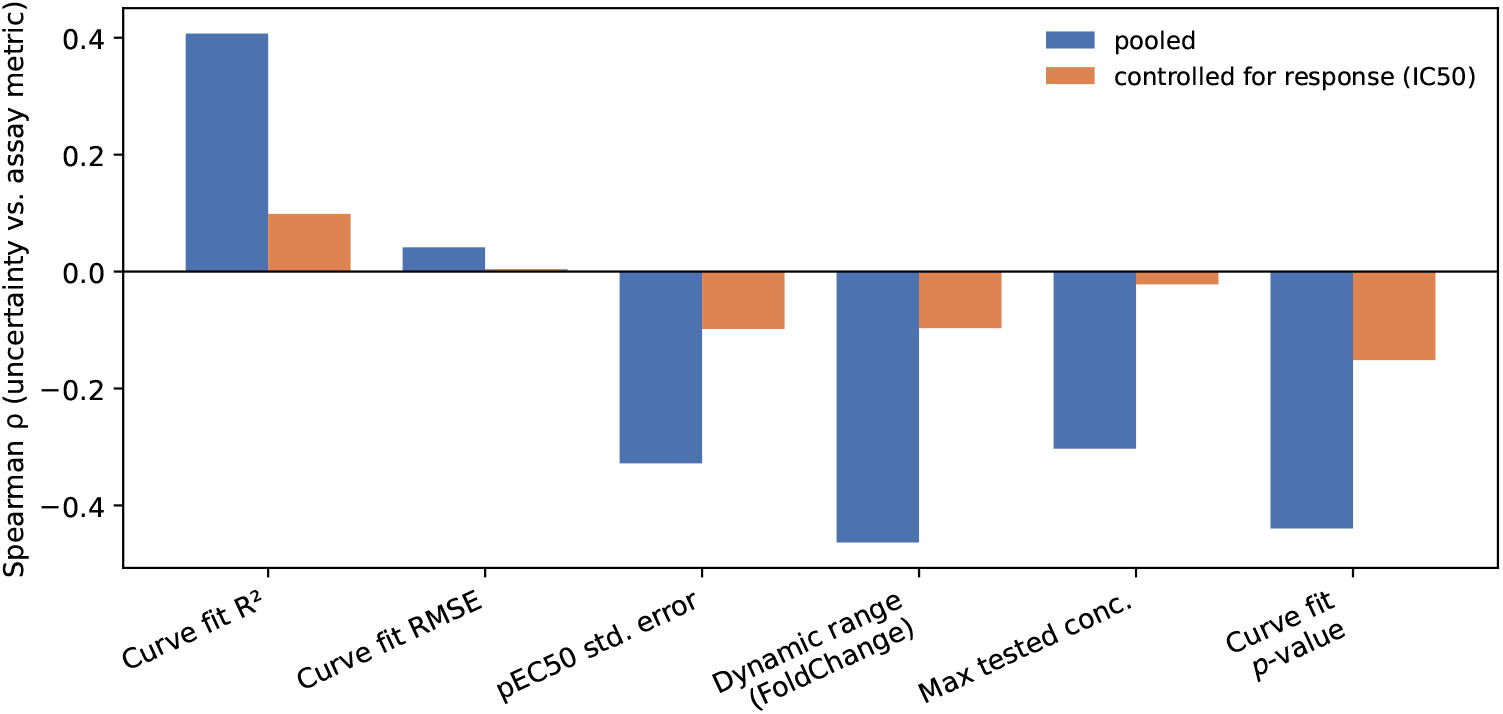
Predicted aleatoric uncertainty of the GaussNNEns on GDSC versus dose-response assay-quality metrics (CurveCurator re-fit of GDSC), shown as Spearman correlations across all pairs (blue) and after controlling for the measured response value (IC50; first-order partial Spearman, orange). The predicted uncertainty correlates moderately with assay properties such as dynamic range and curve-fit *R*^2^, but these associations are largely mediated by the confounding effect of the overall IC50, because the assay metrics are proxies for the response. The root mean squared error (RMSE) of the curve fit is essentially uncorrelated with uncertainty.

**Figure A11.**
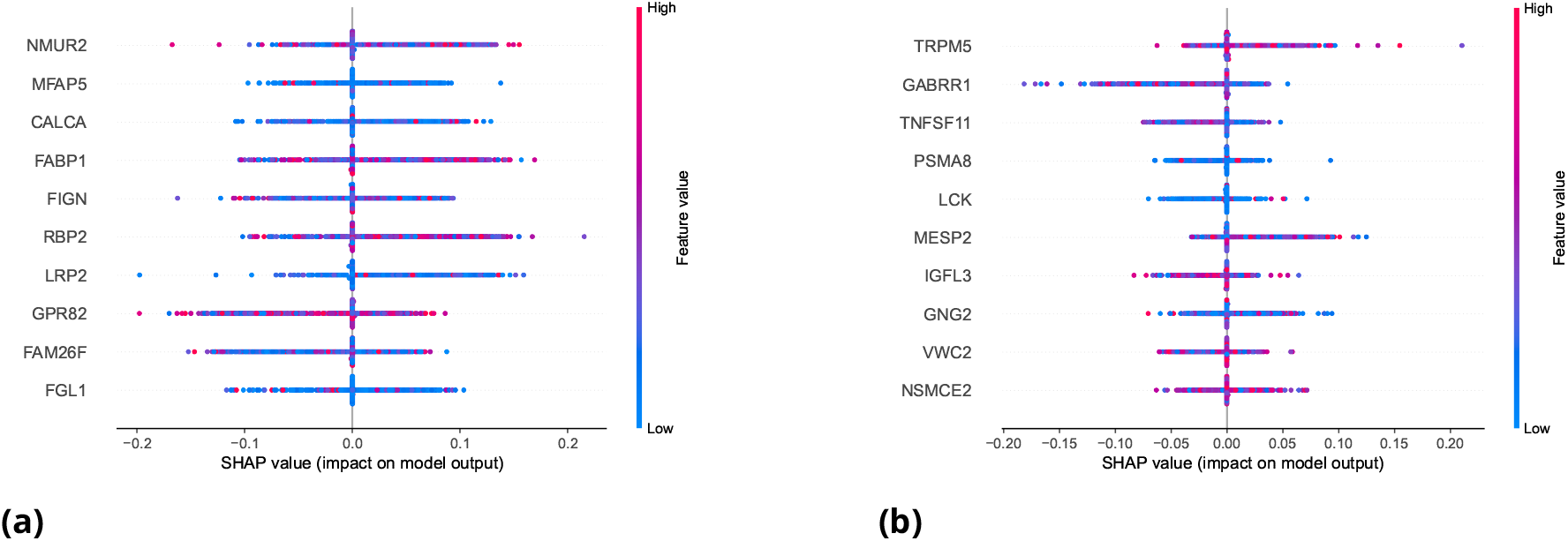
Top-10 gene-level contributions to (a) the predicted mean drug response and (b) the predicted uncertainty of the GaussNN for the test set of one cross-validation fold measured as estimated Shapley values on GDSC. Each point corresponds to a drug-cell-line pair sample. Genes are ordered by overall importance. Notably, distinct sets of genes contribute to response prediction versus uncertainty estimation.

**Figure A12.**
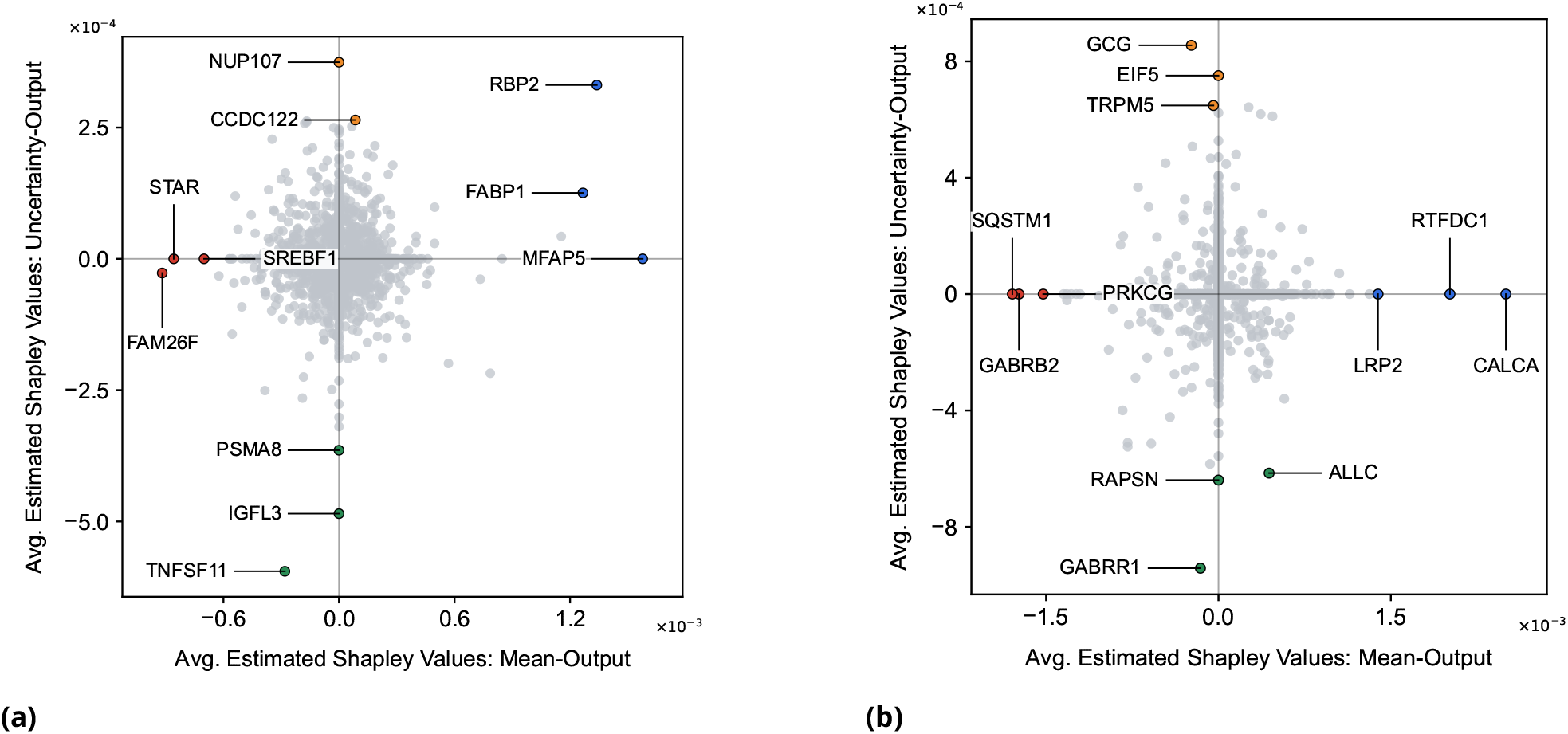
Tissue and pathway stratified estimated Shapley values of gene-expression features for predicted mean drug response and predictive uncertainty for the GaussNN on GDSC for (a) Leukemia and (b) nervous system cell lines with respect to ERK-MAPK targeting drugs. The strongest average contributions across the test set, driving weak response, strong response, uncertainty, and certainty are marked in blue, red, orange, and green, respectively.

**Figure A13.**
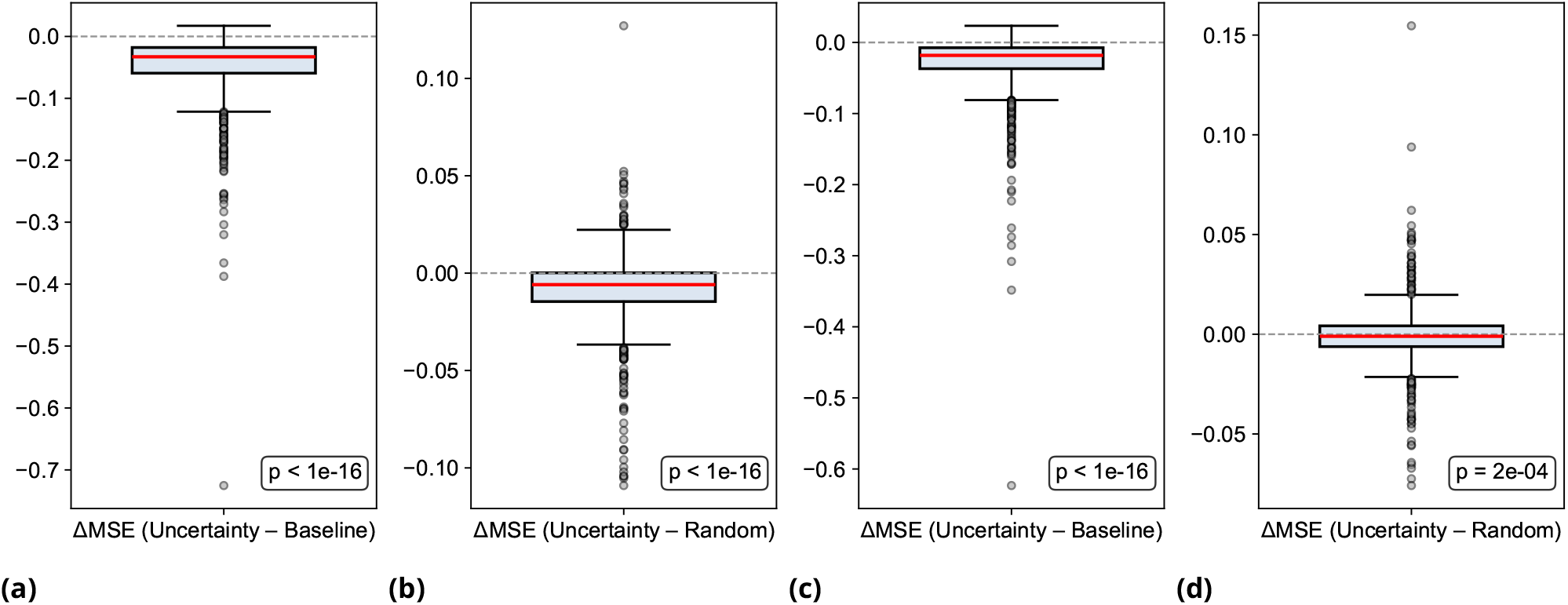
Cell-line–wise differences in mean squared error (ΔMSE) between fine-tuning on an uncertainty-guided, personalized selection of drugs and (a, c) the non-fine-tuned baseline model and (b, d) a model fine-tuned on random drugs, all on GDSC. In (a) and (b), the error is averaged over all drugs; in (c) and (d), it is evaluated only on the subset of drugs not included in either fine-tuning strategy. The GaussNN is first trained on a leave-cell-line-out split, then, for each held-out cell line, fine-tuned with either the 60 drugs with the highest predictive uncertainty or a random subset of 60 drugs added to the training set. Significance is assessed using two-sided Wilcoxon tests.

## Footnotes

1 https://www.rdkit.org/

## References

Abdar M, Pourpanah F, Hussain S, Rezazadegan D, Liu L, Ghavamzadeh M, Fieguth P, Cao X, Khosravi A, Acharya UR, Makarenkov V, Nahavandi S. A review of uncertainty quantification in deep learning: Techniques, applications and challenges. Information Fusion. 2021 Dec; 76:243–297. https://www.sciencedirect.com/science/article/pii/S1566253521001081, doi: 10.1016/j.inffus.2021.05.008.

Adam G, Rampášek L, Safikhani Z, Smirnov P, Haibe-Kains B, Goldenberg A. Machine learning approaches to drug response prediction: challenges and recent progress. npj Precision Oncology. 2020 Jun; 4(1):19. https://www.nature.com/articles/s41698-020-0122-1, doi: 10.1038/s41698-020-0122-1.

Amini A, Schwarting W, Soleimany A, Rus D. Deep Evidential Regression. In: Advances in Neural Information Processing Systems, vol. 33 Curran Associates, Inc.; 2020. p. 14927–14937. https://proceedings.neurips.cc/paper/2020/hash/aab085461de182608ee9f607f3f7d18f-Abstract.html.

Barretina J, Caponigro G, Stransky N, Venkatesan K, Margolin AA, Kim S, Wilson CJ, Lehár J, Kryukov GV, Sonkin D, Reddy A, Liu M, Murray L, Berger MF, Monahan JE, Morais P, Meltzer J, Korejwa A, Jané-Valbuena J, Mapa FA, et al. The Cancer Cell Line Encyclopedia enables predictive modelling of anticancer drug sensitivity. Nature. 2012 Mar; 483(7391):603–607. https://www.nature.com/articles/nature11003, doi: 10.1038/nature11003.

Bernett J, Iversen P, Picciani M, Wilhelm M, Baum K, List M. Critical evaluation of drug response prediction models with DrEval. Nature Communications. 2026 May; 17(1):4238. 10.1038/s41467-026-72903-w, doi: 10.1038/s41467-026-72903-w.

Bishop C. Mixture density networks. Aston University Neural Computing Research Group Report. 1994; .

Chakraborti T, Banerji CRS, Marandon A, Hellon V, Mitra R, Lehmann B, Bräuninger L, McGough S, Turkay C, Frangi AF, Bianconi G, Li W, Rackham O, Parashar D, Harbron C, MacArthur B. Personalized uncertainty quantification in artificial intelligence. Nature Machine Intelligence. 2025 Apr; 7(4):522–530. https://www.nature.com/articles/s42256-025-01024-8, doi: 10.1038/s42256-025-01024-8.

Chung Y, Char I, Guo H, Schneider J, Neiswanger W, Uncertainty Toolbox: an Open-Source Library for Assessing, Visualizing, and Improving Uncertainty Quantification. arXiv; 2021. http://arxiv.org/abs/2109.10254, doi: 10.48550/arXiv.2109.10254, arXiv:2109.10254 [cs].

Corrales E, Levit-Zerdoun E, Metzger P, Mertes R, Lehmann A, Münch J, Lemke S, Kowar S, Boerries M. PI3K/AKT signaling allows for MAPK/ERK pathway independency mediating dedifferentiation-driven treatment resistance in melanoma. Cell Communication and Signaling. 2022 Nov; 20(1):187. 10.1186/s12964-022-00989-y, doi: 10.1186/s12964-022-00989-y.

Fakour F, Mosleh A, Ramezani R, A Structured Review of Literature on Uncertainty in Machine Learning & Deep Learning. arXiv; 2024. http://arxiv.org/abs/2406.00332, doi: 10.48550/arXiv.2406.00332, arXiv:2406.00332 [cs, stat].

Fang Y, Xu P, Yang J, Qin Y. A quantile regression forest based method to predict drug response and assess prediction reliability. PLOS ONE. 2018 May; 13(10):e0205155. https://journals.plos.org/plosone/article?id=10.1371/journal.pone.0205155, doi: 10.1371/journal.pone.0205155.

Gal Y, Ghahramani Z. Dropout as a Bayesian Approximation: Representing Model Uncertainty in Deep Learning. In: Proceedings of The 33rd International Conference on Machine Learning PMLR; 2016. p. 1050–1059. https://proceedings.mlr.press/v48/gal16.html.

Ghandi M, Huang FW, Jané-Valbuena J, Kryukov GV, Lo CC, McDonald ER, Barretina J, Gelfand ET, Bielski CM, Li H, Hu K, Andreev-Drakhlin AY, Kim J, Hess JM, Haas BJ, Aguet F, Weir BA, Rothberg MV, Paolella BR, Lawrence MS, et al. Next-generation characterization of the Cancer Cell Line Encyclopedia. Nature. 2019 May; 569(7757):503–508. 10.1038/s41586-019-1186-3, doi: 10.1038/s41586-019-1186-3.

Gillet JP, Varma S, Gottesman MM. The clinical relevance of cancer cell lines. Journal of the National Cancer Institute. 2013 Apr; 105(7):452–458. doi: 10.1093/jnci/djt007.

Goodfellow IJ, Shlens J, Szegedy C, Explaining and Harnessing Adversarial Examples. arXiv; 2015. http://arxiv.org/abs/1412.6572, doi: 10.48550/arXiv.1412.6572, arXiv:1412.6572 [stat].

Greenman KP, Amini AP, Yang KK. Benchmarking uncertainty quantification for protein engineering. PLOS Computational Biology. 2025 Jul; 21(1):e1012639. https://journals.plos.org/ploscompbiol/article?id=10.1371/journal.pcbi.1012639, doi: 10.1371/journal.pcbi.1012639.

Gruber C, Schenk PO, Schierholz M, Kreuter F, Kauermann G, Sources of Uncertainty in Supervised Machine Learning – A Statisticians’ View. arXiv; 2025. http://arxiv.org/abs/2305.16703, doi: 10.48550/arXiv.2305.16703, arXiv:2305.16703 [stat].

Haibe-Kains B, El-Hachem N, Birkbak NJ, Jin AC, Beck AH, Aerts HJWL, Quackenbush J. Inconsistency in large pharmacogenomic studies. Nature. 2013 Dec; 504(7480):389–393. https://www.nature.com/articles/nature12831, doi: 10.1038/nature12831.

Hirschfeld L, Swanson K, Yang K, Barzilay R, Coley CW. Uncertainty Quantification Using Neural Networks for Molecular Property Prediction. Journal of Chemical Information and Modeling. 2020 Aug; 60(8):3770–3780. 10.1021/acs.jcim.0c00502, doi: 10.1021/acs.jcim.0c00502.

Hu W, Wartmann T, Strecker M, Perrakis A, Croner R, Szallasis A, Shi W, Kahlert UD. Transient receptor potential channels as predictive marker and potential indicator of chemoresistance in colon cancer. Oncology Research. 2023 Nov; 32(1):227–239. https://pmc.ncbi.nlm.nih.gov/articles/PMC10767253/, doi: 10.32604/or.2023.043053.

Iversen P, Witzke S, Baum K, Renard BY. Identifying Drivers of Predictive Aleatoric Uncertainty. In: Proceedings of the Thirty-Fourth International Joint Conference on Artificial Intelligence, vol. 1; 2025. p. 5453–5462. https://www.ijcai.org/proceedings/2025/607, doi: 10.24963/ijcai.2025/607.

Kendall A, Gal Y. What Uncertainties Do We Need in Bayesian Deep Learning for Computer Vision? In: Advances in Neural Information Processing Systems, vol. 30 Curran Associates, Inc.; 2017. https://proceedings.neurips.cc/paper/2017/hash/2650d6089a6d640c5e85b2b88265dc2b-Abstract.html.

Koenker R, Bassett G. Regression Quantiles. Econometrica. 1978; 46(1):33–50. https://www.jstor.org/stable/1913643, doi: 10.2307/1913643.

Kuenzi BM, Park J, Fong SH, Sanchez KS, Lee J, Kreisberg JF, Ma J, Ideker T. Predicting Drug Response and Synergy Using a Deep Learning Model of Human Cancer Cells. Cancer Cell. 2020 Nov; 38(5):672–684.e6. https://www.cell.com/cancer-cell/abstract/S1535-6108(20)30488-8, doi: 10.1016/j.ccell.2020.09.014.

Lakshminarayanan B, Pritzel A, Blundell C. Simple and Scalable Predictive Uncertainty Estimation using Deep Ensembles. In: Advances in Neural Information Processing Systems, vol. 30 Curran Associates, Inc.; 2017. https://papers.nips.cc/paper_files/paper/2017/hash/9ef2ed4b7fd2c810847ffa5fa85bce38-Abstract.html.

Lee BKB, Tiong KH, Chang JK, Liew CS, Abdul Rahman ZA, Tan AC, Khang TF, Cheong SC. DeSigN: connecting gene expression with therapeutics for drug repurposing and development. BMC Genomics. 2017 Jan; 18(1):934. 10.1186/s12864-016-3260-7, doi: 10.1186/s12864-016-3260-7.

Lenhof K, Eckhart L, Rolli LM, Lenhof HP. Trust me if you can: a survey on reliability and interpretability of machine learning approaches for drug sensitivity prediction in cancer. Briefings in Bioinformatics. 2024; 25(5). 10.1093/bib/bbae379, doi: 10.1093/bib/bbae379.

Lenhof K, Eckhart L, Rolli LM, Volkamer A, Lenhof HP. Reliable anti-cancer drug sensitivity prediction and prioritization. Scientific Reports. 2024 May; 14(1):12303. https://www.nature.com/articles/s41598-024-62956-6, doi: 10.1038/s41598-024-62956-6.

MacKay DJC. Bayesian Interpolation. Neural Computation. 1992 May; 4(3):415–447. 10.1162/neco.1992.4.3.415, doi: 10.1162/neco.1992.4.3.415.

Maeda T, Suzuki A, Koga K, Miyamoto C, Maehata Y, Ozawa S, Hata RI, Nagashima Y, Nabeshima K, Miyazaki K, Kato Y. TRPM5 mediates acidic extracellular pH signaling and TRPM5 inhibition reduces spontaneous metastasis in mouse B16-BL6 melanoma cells. Oncotarget. 2017 Sep; 8(45):78312–78326. https://www.oncotarget.com/article/20826/, doi: 10.18632/oncotarget.20826.

Manica M, Oskooei A, Born J, Subramanian V, Sáez-Rodríguez J, Rodríguez Martínez M. Toward Explainable Anticancer Compound Sensitivity Prediction via Multimodal Attention-Based Convolutional Encoders. Molecular Pharmaceutics. 2019 Dec; 16(12):4797–4806. 10.1021/acs.molpharmaceut.9b00520, doi: 10.1021/acs.molpharmaceut.9b00520.

Mentch L, Hooker G. Quantifying Uncertainty in Random Forests via Confidence Intervals and Hypothesis Tests. Journal of Machine Learning Research. 2016; 17(26):1–41. http://jmlr.org/papers/v17/14-168.html.

Nikeghbal P, Zamanian D, Burke D, Steinkamp MP. Organoid models established from primary tumors and patient-derived xenograft tumors reflect platinum sensitivity of ovarian cancer patients. BMC Cancer. 2025 Sep; 25(1):1459. 10.1186/s12885-025-14811-8, doi: 10.1186/s12885-025-14811-8.

Nix DA, Weigend AS. Estimating the mean and variance of the target probability distribution. In: Proceedings of 1994 IEEE International Conference on Neural Networks (ICNN’94), vol. 1; 1994. p. 55–60 vol.1. https://ieeexplore.ieee.org/abstract/document/374138, doi: 10.1109/ICNN.1994.374138.

Nolte D, Ghosh S, Pal R. Efficient Normalized Conformal Prediction and Uncertainty Quantification for Anti-Cancer Drug Sensitivity Prediction with Deep Regression Forests. In: 2024 46th Annual International Conference of the IEEE Engineering in Medicine and Biology Society (EMBC); 2024. p. 1–6. https://ieeexplore.ieee.org/document/10782378, doi: 10.1109/EMBC53108.2024.10782378.

Obermeyer Z, Powers B, Vogeli C, Mullainathan S. Dissecting racial bias in an algorithm used to manage the health of populations. Science. 2019 Oct; 366(6464):447–453. https://www.science.org/doi/10.1126/science.aax2342, doi: 10.1126/science.aax2342.

O’Connor OA, Tobinai K. Putting the Clinical and Biological Heterogeneity of Non-Hodgkin Lymphoma into Context. Clinical Cancer Research. 2014 Oct; 20(20):5173–5181.10.1158/1078-0432.CCR-14-0574, doi: 10.1158/1078-0432.CCR-14-0574.

Padilla OHM, Tansey W, Chen Y. Quantile regression with ReLU Networks: Estimators and minimax rates. Journal of Machine Learning Research. 2022; 23(247):1–42. http://jmlr.org/papers/v23/21-0309.html.

Prasse P, Iversen P, Lienhard M, Thedinga K, Herwig R, Scheffer T. Pre-Training on In Vitro and Fine-Tuning on Patient-Derived Data Improves Deep Neural Networks for Anti-Cancer Drug-Sensitivity Prediction. Cancers. 2022 Jan; 14(16):3950. https://www.mdpi.com/2072-6694/14/16/3950, doi: 10.3390/cancers14163950, number: 16.

Raghavan S. How inclusive are cell lines in preclinical engineered cancer models? Disease Models & Mechanisms. 2022 Jun; 15(5):dmm049520. https://www.ncbi.nlm.nih.gov/pmc/articles/PMC9187871/, doi: 10.1242/dmm.049520.

Rasmussen MH, Duan C, Kulik HJ, Jensen JH. Uncertain of uncertainties? A comparison of uncertainty quantification metrics for chemical data sets. Journal of Cheminformatics. 2023 Dec; 15(1):121. 10.1186/s13321-023-00790-0, doi: 10.1186/s13321-023-00790-0.

Rogers D, Hahn M. Extended-Connectivity Fingerprints. Journal of Chemical Information and Modeling. 2010 May; 50(5):742–754. 10.1021/ci100050t, doi: 10.1021/ci100050t.

Rolli LM, Eckhart L, Herrmann L, Volkamer A, Lenhof HP, Lenhof K. Increasing trustworthiness of machine learning-based drug sensitivity prediction with a multivariate random forest approach. Digital Discovery. 2026; 5(4):1746–1764. 10.1039/d5dd00284b, doi: 10.1039/d5dd00284b.

Seashore-Ludlow B, Rees MG, Cheah JH, Cokol M, Price EV, Coletti ME, Jones V, Bodycombe NE, Soule CK, Gould J, Alexander B, Li A, Montgomery P, Wawer MJ, Kuru N, Kotz JD, Hon CSY, Munoz B, Liefeld T, Dančík V, et al. Harnessing Connectivity in a Large-Scale Small-Molecule Sensitivity Dataset. Cancer discovery. 2015 Nov; 5(11):1210–1223. https://pmc.ncbi.nlm.nih.gov/articles/PMC4631646/, doi: 10.1158/2159-8290.CD-15-0235.

Shi H, Xu T, Li X, Gao Q, Xiong Z, Xia J, Yue Z. DRExplainer: Quantifiable interpretability in drug response prediction with directed graph convolutional network. Artif Intell Med. 2025 May; 163(C). 10.1016/j.artmed.2025.103101, doi: 10.1016/j.artmed.2025.103101.

Peres da Silva R, Suphavilai C, Nagarajan N. TUGDA: task uncertainty guided domain adaptation for robust generalization of cancer drug response prediction from in vitro to in vivo settings. Bioinformatics. 2021; 37(Supplement1):i76–i83. 10.1093/bioinformatics/btab299, doi: 10.1093/bioinformatics/btab299.

Tang YC, Gottlieb A. Explainable drug sensitivity prediction through cancer pathway enrichment. Scientific Reports. 2021 Feb; 11(1):3128. https://www.nature.com/articles/s41598-021-82612-7, doi: 10.1038/s41598-021-82612-7.

Tibshirani RJ, Foygel Barber R, Candes E, Ramdas A. Conformal Prediction Under Covariate Shift. In: Advances in Neural Information Processing Systems, vol. 32 Curran Associates, Inc.; 2019. https://proceedings.neurips.cc/paper/2019/hash/8fb21ee7a2207526da55a679f0332de2-Abstract.html.

Tyner JW, Tognon CE, Bottomly D, Wilmot B, Kurtz SE, Savage SL, Long N, Schultz AR, Traer E, Abel M, Agarwal A, Blucher A, Borate U, Bryant J, Burke R, Carlos A, Carpenter R, Carroll J, Chang BH, Coblentz C, et al. Functional genomic landscape of acute myeloid leukaemia. Nature. 2018 Oct; 562(7728):526–531. https://www.nature.com/articles/s41586-018-0623-z, doi: 10.1038/s41586-018-0623-z.

Wong A, Otles E, Donnelly JP, Krumm A, McCullough J, DeTroyer-Cooley O, Pestrue J, Phillips M, Konye J, Penoza C, Ghous M, Singh K. External Validation of a Widely Implemented Proprietary Sepsis Prediction Model in Hospitalized Patients. JAMA internal medicine. 2021 Aug; 181(8):1065–1070. doi: 10.1001/jamainternmed.2021.2626.

Yang W, Soares J, Greninger P, Edelman EJ, Lightfoot H, Forbes S, Bindal N, Beare D, Smith JA, Thompson IR, Ramaswamy S, Futreal PA, Haber DA, Stratton MR, Benes C, McDermott U, Garnett MJ. Genomics of Drug Sensitivity in Cancer (GDSC): a resource for therapeutic biomarker discovery in cancer cells. Nucleic Acids Research. 2013 Jan; 41(D1):D955–D961. 10.1093/nar/gks1111, doi: 10.1093/nar/gks1111.

Zech JR, Badgeley MA, Liu M, Costa AB, Titano JJ, Oermann EK. Variable generalization performance of a deep learning model to detect pneumonia in chest radiographs: A cross-sectional study. PLOS Medicine. 2018 Nov; 15(11):e1002683. https://journals.plos.org/plosmedicine/article?id=10.1371/journal.pmed.1002683, doi: 10.1371/journal.pmed.1002683.

